# Mechanical competition triggered by innate immune signaling drives the collective extrusion of bacterially-infected epithelial cells

**DOI:** 10.1101/2020.01.22.915140

**Authors:** Effie E. Bastounis, Francisco Serrano Alcalde, Prathima Radhakrishnan, Patrik Engström, María J. Gómez Benito, Mackenzi S. Oswald, Jason G. Smith, Matthew D. Welch, José M. García Aznar, Julie A. Theriot

## Abstract

Multiple distinct types of intracellular bacterial pathogens have been shown to alter the mechanics of their mammalian host cells to promote cell-to-cell spread. Conversely, host cells may respond by altering their own mechanical behavior to limit infection. We monitored epithelial cell monolayers sparsely infected with the intracellular bacterial pathogens *Listeria monocytogenes* or *Rickettsia parkeri* over the course of several days. Under conditions where these pathogens were able to trigger innate immune signaling through the NF-κB pathway and to use actin-based motility to spread non-lytically from cell to cell, domains of infected cells formed enormous three-dimensional mounds, where uninfected cells surrounding the infected cells became stiffer and actively moved toward the site of infection, collectively squeezing the softer and weaker infected cells up and ejecting them from the epithelial monolayer. Bacteria in mounds were less able to spread laterally in the monolayer, limiting the growth of the focus, while mounded cells eventually underwent cell death. Cells in infected monolayers exhibited behavioral and molecular signatures of the epithelial to mesenchymal transition (EMT), such that coordinated forceful action by uninfected bystander cells actively eliminated large domains of infected cells, consistent with the hypothesis that this collective cell response represents an innate immune response.

## INTRODUCTION

During bacterial infection, host cells and pathogens communicate using a wide variety of chemical and biochemical signals. Also important, but sometimes underappreciated, are mechanical signals that contribute to host-pathogen interactions. Example of mechanical signals include the forces that cells are able to transduce to each other and to their extracellular matrix. These cell-generated forces are critical for maintaining tissue integrity and barrier functions.

The biomechanical interactions between host cells and bacterial pathogens are dictated by both the pathogen and the host. It has been previously shown that intracellular pathogens can manipulate host cell functions, including host cell mechanotransduction, in order to spread more efficiently (Faralla et al., 2018; Lamason et al., 2016; Rajabian et al., 2009). For example, two intracellular bacterial pathogens that generate actin comet tails for propulsion through the cytoplasm of infected host epithelial cells, *Listeria monocytogenes* (*L.m.*) and *Rickettsia parkeri* (*R.p.*), both secrete virulence factors that reduce the tension at cell-cell junctions, making it easier for the bacteria to propel themselves into neighboring cells to spread the infection. *R.p.* secretes Sca4, which decreases intercellular tension at apical junctions in polarized epithelial cells by binding to vinculin and inhibiting its association with α-catenin (Lamason et al., 2016). *L.m.* secretes an unrelated virulence factor, InlC, which reduces tension at cell-cell junctions by a distinct biochemical mechanism, binding and inhibiting the activity of the mammalian adaptor protein Tuba (Rajabian et al., 2009). In addition, *L.m.* secretes an additional factor, InlP, that binds to yet another host target protein, afadin, leading to a weakening of the tensional forces between host epithelial cells and their basement membrane extracellular matrix, thus facilitating apico-basal bacterial spread in the placenta (Faralla et al., 2018).

Conversely, epithelial cells may exhibit biomechanical alterations after bacterial infection that appear to function as innate immune responses limiting bacterial spread. For example, in intestinal epithelial cells infected with *Salmonella enterica* serovar Typhimurium, (*S.Tm.*) innate immune activation of the inflammasome triggers apical extrusion of individual infected cells, which are then shed into the intestinal lumen, while the integrity of the epithelial barrier is maintained (Knodler et al., 2014; Sellin et al., 2014). Similar extrusion of single infected cells has also been observed in human intestinal enteroids infected with *S.Tm*. or *L.m.* (Co et al., 2019). The biomechanical mechanism leading to extrusion of individual infected cells from polarized epithelia is similar to the mechanism that normal, uninfected epithelia use to extrude dying apoptotic cells, or excess cells in the context of overcrowding (Gudipaty and Rosenblatt, 2017). Typically, a dying cell begins to contract, and signals its immediate neighbors to assemble a multicellular contractile actomyosin “purse-string” that contracts and moves basally, squeezing out the extruding cell while promoting formation of new cell-cell junctions among the new neighbors (Kuipers et al., 2014; Rosenblatt et al., 2001; Tamada et al., 2007). At low epithelial cell density, active lamellipodial protrusion and crawling of the neighboring cells also contributes to apoptotic cell extrusion (Kocgozlu et al., 2016). Some bacterial pathogens resist this innate immune extrusion reaction by secreting virulence factors that specifically stabilize cell-matrix adhesions (Kim et al., 2009).

A particular challenge is faced by epithelia that are infected by pathogens capable of direct, non-lytic cell-to-cell spread. In addition to *L.m.* and *R.p.*, a wide variety of other unrelated intracellular bacterial pathogens have independently developed biochemically distinct mechanisms to trigger the rapid assembly of host cell actin filaments at one pole of the bacterial cell (Stevens et al., 2006). The substantial pushing forces generated by actin network assembly propel the bacteria rapidly through host cell cytoplasm and into membrane-bound protrusions at the host cell surface (Tilney and Portnoy, 1989). In the context of an epithelial monolayer, these protrusions facilitate direct cell-to-cell spread of bacteria into neighboring cells, without causing lysis of the originally infected host cell (Lamason and Welch, 2017). Efficient actin-driven cell-to-cell spread of *L.m.* in cultured epithelial cells can give rise to large infected domains or foci comprising hundreds of epithelial cells in less than 24 h after the initial invasion of a single cell (Ortega et al., 2019). In this situation, extrusion of individual infected cells using a purse-string mechanism would clearly be insufficient to limit the infection.

Infection of epithelial cells by many bacterial pathogens triggers activation of the transcription factor NF-κB (Dev et al., 2011). In normal cells at rest, NF-κB is inactive and retained in the cytoplasm. During infection, pathogen-associated molecular patterns can initiate a signal transduction cascade that leads to activation of NF-κB and its translocation into the nucleus, where it can induce the expression of additional genes encoding cytokines and adhesion molecules, promoting the recruitment of immune cells to the site of infection (Jiang et al., 2011). For several intracellular bacterial pathogens, including *L.m.*, infection of epithelial cells in a monolayer triggers activation of NF-κB and subsequent cytokine production not only in the infected cells themselves, but also in nearby uninfected bystander cells (Kasper et al., 2010). This suggests that uninfected bystander cells may play a role in innate immune response to infection by amplifying the alarm signals produced by infected cells.

To determine whether and how epithelial monolayers might alter their biomechanical behavior to limit the spread of pathogens capable of direct, non-lytic cell-to-cell spread, we used long-term time-lapse videomicroscopy to monitor individual infected foci of cultured epithelial monolayers for several days after exposure to *L.m.* or *R.p*. For both pathogens, we found that the uninfected neighboring cells undergo a dramatic behavioral change with similarities to the epithelial to mesenchymal transition (EMT), and act collectively to squeeze and eventually force the extrusion of large domains of infected cells, forming enormous “mounds”. Mound formation is driven by changes in cellular stiffness and force transduction in both of the two battling cell populations, the uninfected “surrounders” and the infected “mounders”, and critically depends on cell-cell force transduction through adherens junctions. Innate immune signaling pathways, and specifically NF-κΒ activation in both the infected cells and in uninfected bystanders, are drivers in this mechanical competition. Importantly, we found that wild-type *R.p.*, which actively suppresses NF-κB activation in infected host cells, do not induce mound formation, while *ompB-R.p.* triggers both NF-κB activation and mound formation. This result demonstrates that the mechanical changes in epithelial cells leading to mound formation are not simply a result of cytoskeletal remodeling associated with bacterial infection and cell-to-cell spread, but rather are specifically triggered by innate immune signaling and probably serve a protective function for the host. Furthermore, we find that both infected cells and uninfected bystander cells (“surrounders”) upregulate molecular markers associated with the EMT, as well as exhibit EMT-like mechanical and behavioral changes. This indicates that uninfected bystander cells can contribute to the innate immune response to infection not only by amplifying the chemical signals that recruit immune cells, but also by a collective mechanical ejection of the infected domain. Overall, our findings connect the EMT, a widely conserved cell behavioral transformation which is critical in many stages of embryonic development as well as pathologies including fibrosis and cancer, with innate immune mechanisms. These results also underline the dynamic remodeling capability of epithelial tissue, and lead us to propose that coordinated mechanical forces may be a mechanism employed by host epithelial cells to eliminate bacterial infection.

## RESULTS

### Infection with *Listeria monocytogenes* (*L.m.*) alters the organization and kinematics of host epithelial cells to form large multicellular mounds

To address whether infection elicits changes in the kinematic behavior of host cells in epithelial monolayers, we used Madin-Darby Canine Kidney (MDCK) epithelial cells as model host cells because they form well-polarized and homogeneous monolayers in culture and have often been used to study *L.m.* infection (Ortega et al., 2019; Pentecost et al., 2006; Robbins et al., 1999). MDCK cells grown in a confluent monolayer on glass coverslips were infected with low dosage of wild-type *L.m*. so that on average fewer than one in 10^3^ host cells were invaded, resulting in well-spaced infectious foci that could be observed over a period of several days. Cells were maintained with the membrane-impermeable antibiotic gentamicin in the culture medium so that bacteria could only spread directly from cell to cell, and the *L.m.*-infected foci grew to include several hundred host cells over a period of 24-48 h. Whereas MDCK cells in uninfected monolayers exhibited regularly spaced nuclei confined to a single plane (Figure 1A), cells in *L.m.*-infected foci formed large three-dimensional mounds over the time of observation, with host cells piling on top of one another to form structures up to 50 µm tall (Figure 1B and Figure S1A-C), with an average volume of extruded cells ∼1.7 x 10^5^ µm^3^ after 24 h (N=77 mounds). We observed formation of morphologically similar but larger infection mounds in monolayers of human A431D epithelial cells expressing normal human E-cadherin (average volume at 24 h ∼1.3 x 10^6^ µm^3^, N=5), indicating that this phenomenon is not unique to canine epithelial cells (Figure S1D). We also infected monolayers of normal primary human enteroid-derived cells obtained from ileal resection, and observed prominent mound formation (Figure S1E-F, average volume at 24 h ∼2.6 x 10^5^ µm^3^, N=9). These results suggest that mound formation may be a widely conserved phenomenon for mammalian epithelial monolayers, including both immortalized cell lines and untransformed primary cells.

**Figure 1.**
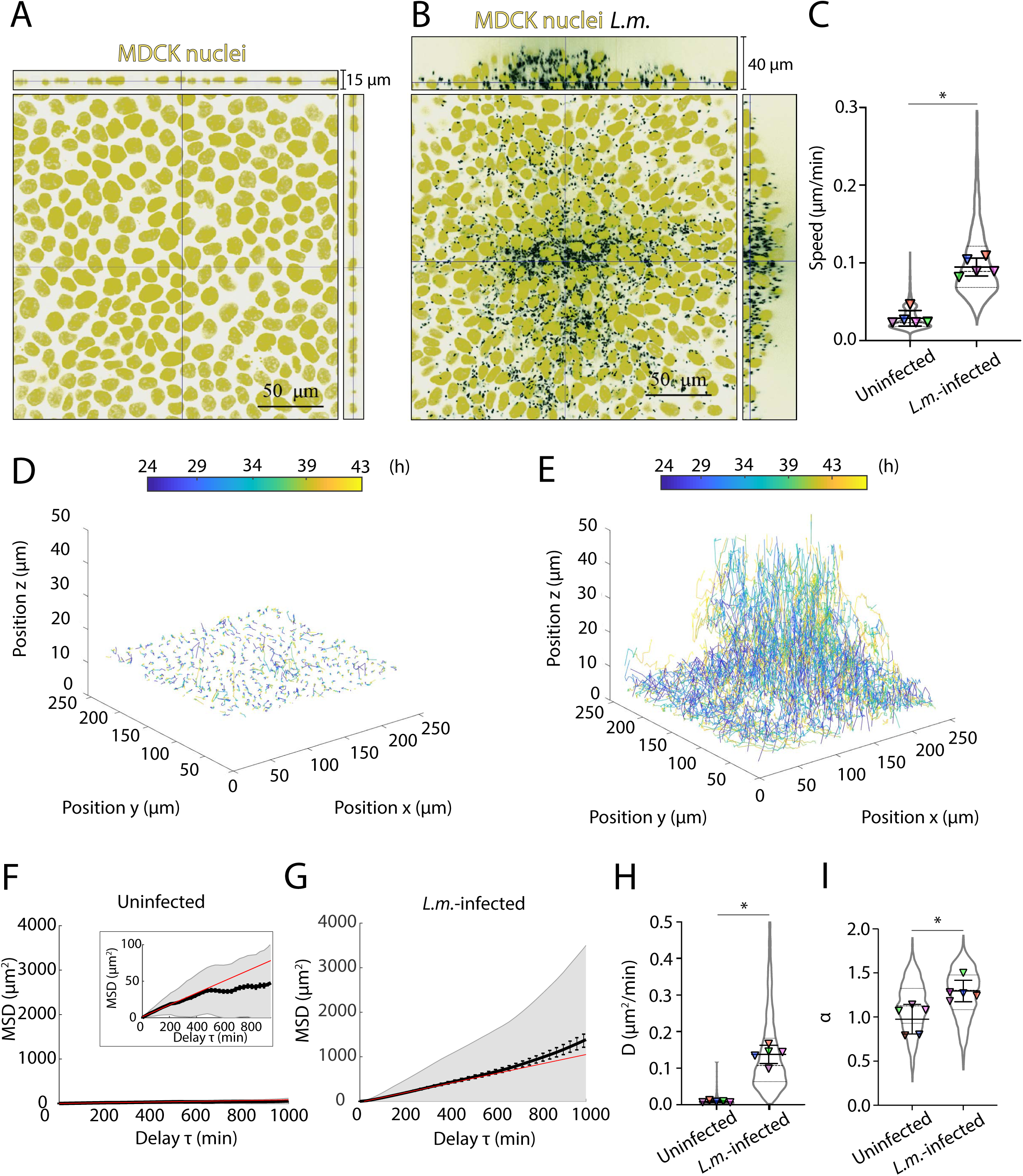
Kinematics of epithelial cell monolayers infected with *Listeria monocytogenes*. (*L.m.*) change dramatically 24 p.i.. (**A-B**) Representative orthogonal views of (A) uninfected and (B) *L.m.*-infected MDCK cells in a confluent monolayer (host nuclei: yellow, *L.m.*: black) at 24 h p.i.. For (B) a single infection mound is shown. (**C**) Violin plots of mean speed of uninfected and *L.m.*-infected MDCK nuclei tracked between 24 to 44 h p.i.. (**D-E**) 3D displacements of (D) uninfected and (E) *L.m.*-infected MDCK nuclei tracked between 24 to 44 h p.i.. Nuclear trajectories are colored based on time. (**F-G**) Plots of the weighted average for each time delay of all mean squared displacement (MSD) curves of MDCK nuclei in a particular field of view tracked between 24 to 44 h p.i.. Panels F-G refer to uninfected and *L.m.*-infected MDCK respectively (grayed area: weighted standard deviation over all MSD curves; error bars: standard error of the mean, see STAR methods). Red line is the weighted mean MSD curve from which the diffusion coefficient *D* is calculated. The fit was made in the first 500 min. In panel F a zoomed inset is provided. **(H-I**) Violin plots of average diffusion coefficient *D* (H) and scaling exponent *α* (I) for uninfected and *L.m.*-infected MDCK nuclei tracked between 24 to 44 h p.i. (n=1259 and n=1397 nuclei respectively). For (C) and (H-I) dashed line in the violin plots is the median and dotted lines the 25 and 75% quartiles. For each experiment (N=5) the average value over the entire field of view and over time is shown as a triangle (mean+/-SD, unpaired T-test: * p<0.01). See also Figure S1 and Movie S1.

Formation of these large mounds was due to active directional movement by the host epithelial cells. From 24 h post-infection (p.i.) onwards, we monitored the movement of Hoechst-stained host cell nuclei via time-lapse confocal 3D microscopy. Uninfected MDCK cells in confluent monolayers moved randomly with very low speeds and rarely translocated more than a fraction of one cell diameter over 20 h, appearing to be confined by their neighbors (Movie S1; Figure 1C-D). In contrast, *L.m.*-infected cells within foci and their nearby uninfected neighbors all moved rapidly and persistently toward the center of the focus, and the infected cells also moved upward, away from the coverslip, to form the three-dimensional mound (Movie S1; Figure 1C, E). Net cell speeds inside and near *L.m.*-infected foci increased by more than 3-fold relative to uninfected monolayers (Figure 1C), and cell tracks became significantly straighter (Figure S1G). The dramatic nature of this change in cell behavior is better appreciated by examination of the mean squared displacement (MSD) for cells in uninfected versus infected monolayers. Over 16 h of observation, the average MSD for uninfected cells remained under 50 µm^2^, indicating a net displacement of about 7 µm. Movement over this time frame was strongly constrained, with the average MSD reaching a plateau, rather than continuing to grow linearly with time as would be expected for unconstrained random movement (Figure 1F). In contrast, the average MSD for cells in and near *L.m.*-infected foci grew to over 1000 µm^2^ over this same time frame, and increased superdiffusively, faster than linear with time, indicative of directed motion (Figure 1G-I). Altogether these findings indicate that the host cells undergo a behavioral phase transition during infection, switching from a caged state where they are tightly confined by their neighbors to a superdiffusive, highly motile state at late times p.i.. Such transitions have been previously described for epithelia undergoing the epithelial to mesenchymal transition (EMT) or for cells exposed to compressive forces (Park et al., 2016) but not for cells during bacterial infection.

These behavioral changes were accompanied by changes in cell size, shape and packing in the epithelial monolayer. To measure changes in cell morphology, we performed cell segmentation based on the pericellular localization of the adherens junction protein E-cadherin (Figure S2A). We used the fluorescent signal from *L.m.* constitutively expressing red fluorescent protein to identify the cells containing the bacteria forming the mound (“infected mounders”) and their nearby uninfected neighbors (“surrounders”). In uninfected monolayers, the MDCK cells formed regularly packed polygons, with the long axis of each cell oriented randomly with respect to the center of the field of view (Figure 2A). At foci in infected monolayers, infected mounders at the center of the focus had smaller projected 2D area than uninfected cells or uninfected surrounders (Figure 2A-B). Strikingly, the orientation of the long axis of the surrounders pointed toward the center of the focus, and this preferential orientation persisted over more than 100 µm from the site of the mound, equivalent to more than 10 cell diameters (Figure 2A, C). These surrounder cells also showed significantly more alignment of their long axes with respect to their immediate neighbors, indicating an increase in local nematic order within the monolayer (Figure S2B). In addition we calculated the aspect ratio *AR* and shape factor *q* (ratio of cell perimeter to square root of area), dimensionless numbers that have previously been used to characterize cell shape changes during EMT (Mitchel et al., 2019), and found increases for both values in both infected mounders and surrounders as compared to cells in uninfected monolayers (Figure S2C-D).

**Figure 2.**
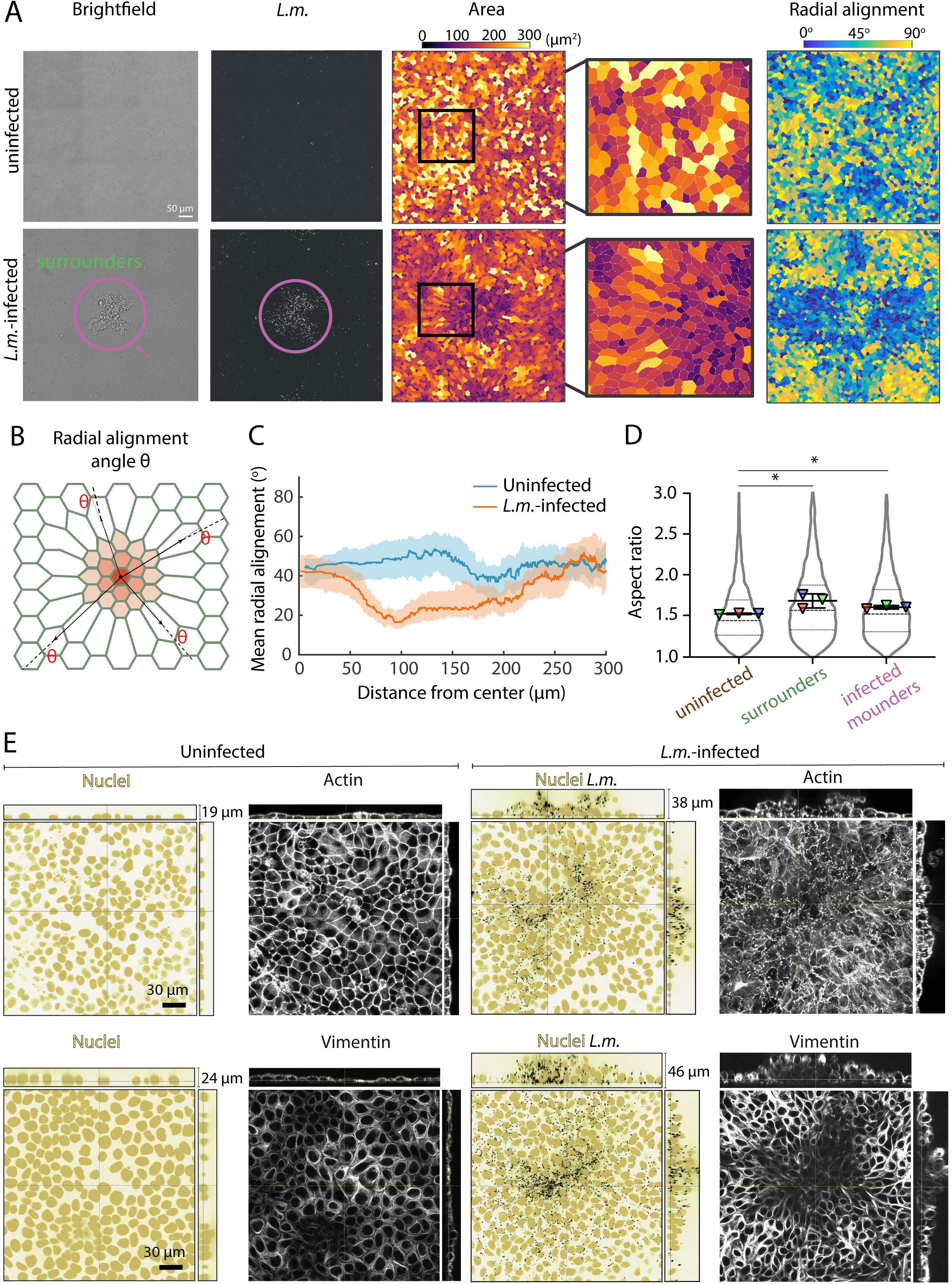
*L.m.*-infected MDCK cells alter their shape, orientation and cytoskeletal organization late in infection. **(A)** Changes in cell area and orientation for uninfected (top) and *L.m.*-infected MDCK cells (bottom) at 24 h p.i.. Columns: brightfield image, corresponding *L.m.* fluorescence, segmented cells color-coded based on their area and on their radial alignment angle (see STAR methods). Purple circle points on an infection mound. 4^th^ column shows a zoomed portion of the area of cells adjacent to the edge of the mound. **(Β**) Schematic showing how radial alignment is defined as the angle θ between the radial direction and the direction of the major axis of the cell. If the orientation is close to zero, cells are oriented radially towards the centre of the infection focus. If the orientation is close to 90° the orientation is circumferential. **(C**) Radial alignment angle θ (x-axis) versus distance from the center of the field of view (y-axis) for uninfected (blue) and *L.m.*-infected cells (orange) shown in panel A. Solid line shows the median and shaded areas the 40 and 60% percentiles. **(D**) Violin plot of aspect ratio (major/minor axis) for uninfected cells (n=5817), surrounders (n=5163) and mounders (n=1248). Dashed line: median, dotted lines: 25 and 75% quartiles. For each experiment (N=3) the average value over the entire field of view is shown as a triangle (mean+/-SD, unpaired t-test: * p<0.05). **(E)** Representative orthogonal views of uninfected (1^st^ column) or *L.m.*-infected MDCK monolayers (3^rd^ column) (host nuclei: yellow, *L.m.*: black) and corresponding F-actin (2^nd^ and 4^th^ column, top) and vimentin localization (2^nd^ and 4^th^ column, bottom). See also Figure S2.

The changes in cell morphology for both infected mounders and surrounders suggested that there might be cytoskeletal rearrangements associated with infection. We performed immunofluorescence on *L.m.-*infected samples at 24 h p.i. and examined them using confocal microscopy. We found that F-actin localized pericellularly for cells in uninfected wells, whereas in infected wells both infected mounders and surrounders contained far less pericellular actin, while actin comet tails were readily visible in infected cells (Figure 2D). In addition, surrounders exhibited prominent F-actin stress fibers compared to infected mounders and to uninfected cells. Microtubules were also disrupted in infected cells, as were the intermediate filaments vimentin and cytokeratin, with vimentin most strongly reduced in the infected mounders relative to uninfected surrounders (Figure 2D and S2E-F). Altogether, the changes in the cytoskeletal organization of infected mounders as compared to surrounders is consistent with these cells displaying distinct mechanical properties that could contribute to the formation of the 3D mounds. Alternatively, changes in the organization of cytoskeletal filaments could be the result of infected mounders being squeezed and/or extruded rather than the initial cause of mounding.

### Mechanical cellular competition drives infected cell extrusion en masse

We found that infected mounders and surrounders have distinct shape characteristics and cytoskeletal organization when compared to each other and to uninfected cells. This led us to hypothesize that the formation of an extruded mound might require competition between two cell populations with distinct biomechanical properties. Indeed, when we infected epithelial cells with a high multiplicity of infection (MOI) so that all cells became infected, we did not observe mound formation (Figure 3A-B). This finding suggests that mound formation may not simply be a consequence of loss of cell-substrate adhesion by infected cells, but instead may require the involvement of uninfected surrounder cells in a form of mechanical competition.

**Figure 3.**
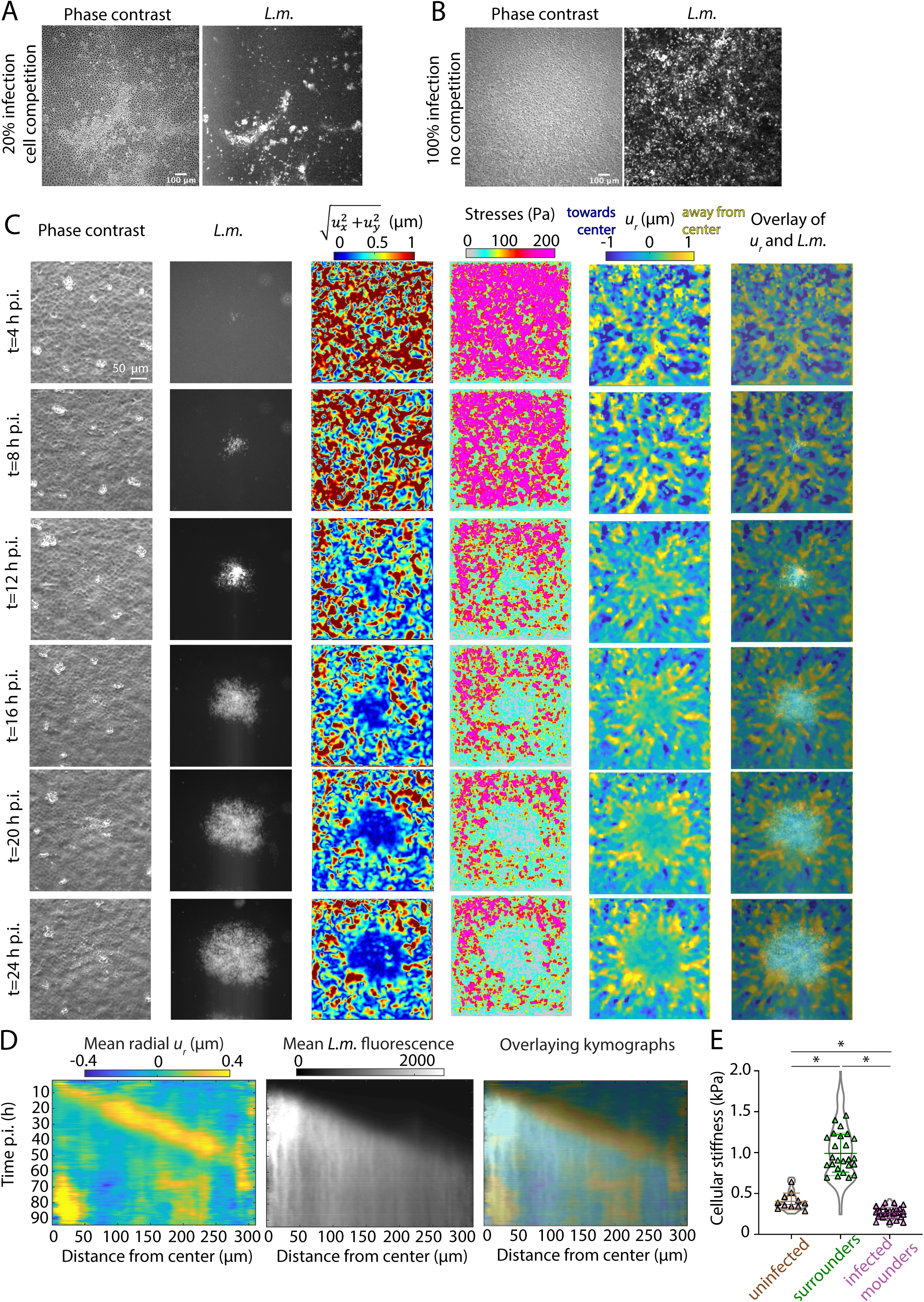
Competition between surrounders and infected mounders is mechanically driven. **(A)** Phase contrast image of MDCK cells (left) and corresponding *L.m.* fluorescence (right) for cells that are 20% infected at 24 h p.i.. An infection mound can be observed in the phase contrast image. **(B)** Same as panel A but for cells that are 100% infected. Lack of cell competition results in lack of infection mounding. **(C)** Representative phase contrast images of confluent MDCK cells adherent on 3 kPa gels and infected with low multiplicity of *L.m.*, corresponding *L.m.* fluorescence, cell-matrix deformation maps (color indicates deformation magnitude in µm), traction stresses exerted by cells on their matrix (color indicates stress magnitude in Pa), radial cell-matrix deformation, *u_r_* maps (positive deformations values in µm indicate deformations pointing away from the center of the focus, outwards) and overlap of *u_r_* maps and *L.m.* fluorescence. Rows point to different times p.i.. (**D**) Kymograph of the mean radial deformation *u_r_* (in µm) as a function of time, t (left). The center of the polar coordinate system is considered at the center of the infection focus. Kymograph of the mean radial *L.m.* fluorescence as a function of time, t (middle). Overlay of both previous kymographs (right). **(E)** Violin plots of measurements of cellular stiffness (kPa) for cells originating from uninfected wells (n=68), surrounders (n=126) and infected mounders (n=130) indented 2-8 times. Dashed line: median, dotted lines: 25 and 75% quartiles. For each cell its average stiffness is shown as a triangle (mean+/-SD, Wilcoxon Rank Sum test: * p<0.01). See also Figure S3 and Movie S2.

To test whether there is a difference in the mechanical properties of the surrounders versus the infected mounders, we first used traction force microscopy (TFM) to measure cellular traction forces exerted by cells on the extracellular matrix on which they reside. In TFM, cells actively pull on their matrix depending on how well their focal adhesions are organized and connected to the underlying cytoskeleton, and cellular force generation can be inferred from displacement of fluorescent tracer particles embedded in the deformable matrix (Bastounis et al., 2014; del Álamo et al., 2007). To that end, we placed MDCK cells in confluent monolayers on soft 3 kPa collagen I-coated hydrogels and subsequently infected them with *L.m.*. We found that, as the infection focus grew, the cells within the mound progressively adhered more weakly to their matrix as compared to surrounding uninfected cells, as evidenced by reduced total deformation and significantly reduced traction stresses directly underneath the infected cells in each focus (Figure 3C; Movie S2). Importantly, when we converted the deformations into polar coordinates and examined the radial deformations *u_r_*, we found that the surrounders just at the edge of the infection mound exerted particularly large deformations, grabbing the substrate and pulling it outwards (*i.e.* away from the mound) as they moved directionally towards the center of the focus (Figure 3C). This orientation of traction force generation by the surrounders is not consistent with extrusion generated by a contractile purse-string mechanism, but is consistent with lamellipodial protrusion and directed cell migration on the part of the surrounders (Kocgozlu et al., 2016). Overall these dynamic changes in traction force generation patterns indicate that mound formation is not caused by contraction of the infected cells, but rather by active crawling movement of the uninfected surrounders, forcefully engaging the substrate to migrate toward the center of the focus, squeezing and extruding the domain of infected cells.

When we constructed kymographs by calculating the average radial deformation and *L.m.* fluorescence intensity as a function of radial distance from the center of the mound, we noted that the largest radial deformations *u_r_* of the substrate coincided with the edge of the infection focus, indicating that the uninfected surrounders immediately adjacent to the infected focus generated the strongest outward-directed forces (Figure 3D, Figure S3A-B). This behavior persisted up to 54 h p.i. with the peak of forces moving outwards as the infection focus grew. However, after 54 h p.i. the peak bacterial fluorescence started decreasing, presumably due to infected cells having been extruded from the monolayer and bacteria being therefore unable to spread anymore to new uninfected cells at the basal cell layer. Concurrently, the traction forces underneath the infectious focus began to rise, suggestive of uninfected cells repopulating the region where the weakly adhering infected mounders had previously resided (Movie S2).

The decrease in the traction stresses of the infected mounders could be caused by the activity of secreted *L.m.* virulence factors previously demonstrated to weaken cellular tension by interacting with either cell-cell or cell-matrix adhesion complexes, such as the internalins C (InlC) (Rajabian et al., 2009) and P (InlP) (Faralla et al., 2018). To test this possibility, we infected MDCK cells with *ΔinlC^-^* or *ΔinlP^-^* mutants and assessed the ability of host cells to form infection mounds at 24 h p.i.. Since the pore-forming toxin listeriolysin O (LLO) could also play a role by activating signaling pathways that could affect mechanotransduction, we also infected cells with LLO^G486D^ *L.m.* a strain carrying a point mutation in the gene encoding LLO that exhibits 100x less hemolytic activity than the wild-type protein, permitting a moderate degree of bacterial proliferation and cell-to-cell spread (unlike strains completely lacking LLO (*Δhly^-^*) which are unable to proliferate in host cells) (Rengarajan et al., 2016). In all cases we were able to observe infection mounds at 24 h p.i.. While we found no differences in mound volume when comparing *ΔinlC^-^* and *ΔinlP^-^* to wild-type (WT) *L.m.* infection mounds, we did find that infection with LLO^G486D^ *L.m.* led to formation of smaller infection mounds compared to the other conditions (Figure S3C-D). However, it is worth noting that the total bacterial load in the LLO^G486D^ *L.m.* mounds was also decreased, to an extent comparable to the degree of reduction of the volume of the mound (Figure S3E). Altogether, these results suggest that neither InlC nor InlP secreted bacterial virulence factors contribute to infection mound formation, reinforcing the idea that this cellular competition leading to infection mounding might be determined by the host cells themselves rather than by the bacteria.

The decrease in the traction forces that infected mounders exerted on their substrate could be due to specific changes in the organization of their focal adhesions or to changes in the cytoskeletal integrity of cells, resulting in reduced ability of the cells to transmit forces. To determine whether the bulk cytoskeletal stiffness of cells changes upon infection, we used atomic force microscopy (AFM) to measure the stiffness of surrounders or infected mounders at 24 h p.i. as well as the stiffness of uninfected cells in MDCK monolayers that had never been exposed to *L.m*.. We found that stiffness of surrounders was on average 4-fold higher than of infected mounders. As compared to cells originating from uninfected wells, surrounders were 2.5-fold stiffer, whereas infected mounders were 1.6-fold softer (Figure 3E), consistent with the cytoskeletal disruption of the infected cells as previously noted in Figure 2D.

Combined with the TFM results described above, these measurements help us to describe the sequence of mechanical events that results in mound formation. As compared to the steady-state cell behavior in uninfected epithelial monolayers, infected cells become softer, and lose the ability to generate strong traction forces on their substrates. At the same time, the nearby uninfected surrounder cells, particularly those physically closest to the infected cells, become stiffer and directionally polarized, gripping the substrate and actively pulling on it to move themselves directionally and persistently toward the center of the focus. The mechanical competition between the surrounders and the infected mounders results in the mounders getting squeezed and eventually ejected from the plane of the monolayer.

### Cell-cell adhesions and actomyosin contractility contribute to infection mound formation

As infected mounders and surrounders differ in cellular stiffness, cytoskeletal organization, and ability to generate traction force on their substrates, we hypothesized that perturbation of cell-matrix or cell-cell force transduction via pharmacological or genetic manipulation should inhibit the collective extrusion of infected cells and reduce mound formation. We first tested how inhibition of cell-matrix contractility would affect infection mounding by infecting MDCK monolayers with *L.m.* and treating cells 4 h p.i. with the ROCK inhibitor Y-27632 or the myosin II inhibitor blebbistatin to decrease actomyosin-generated contractility. In both cases we found a significant reduction in infection mound volume relative to vehicle control at 24 h p.i. (Figure 4A-B). This result reinforces the idea that cell-matrix contractility and traction force generation by the surrounders is crucial to eliciting the squeezing and subsequent extrusion of infected mounders.

**Figure 4.**
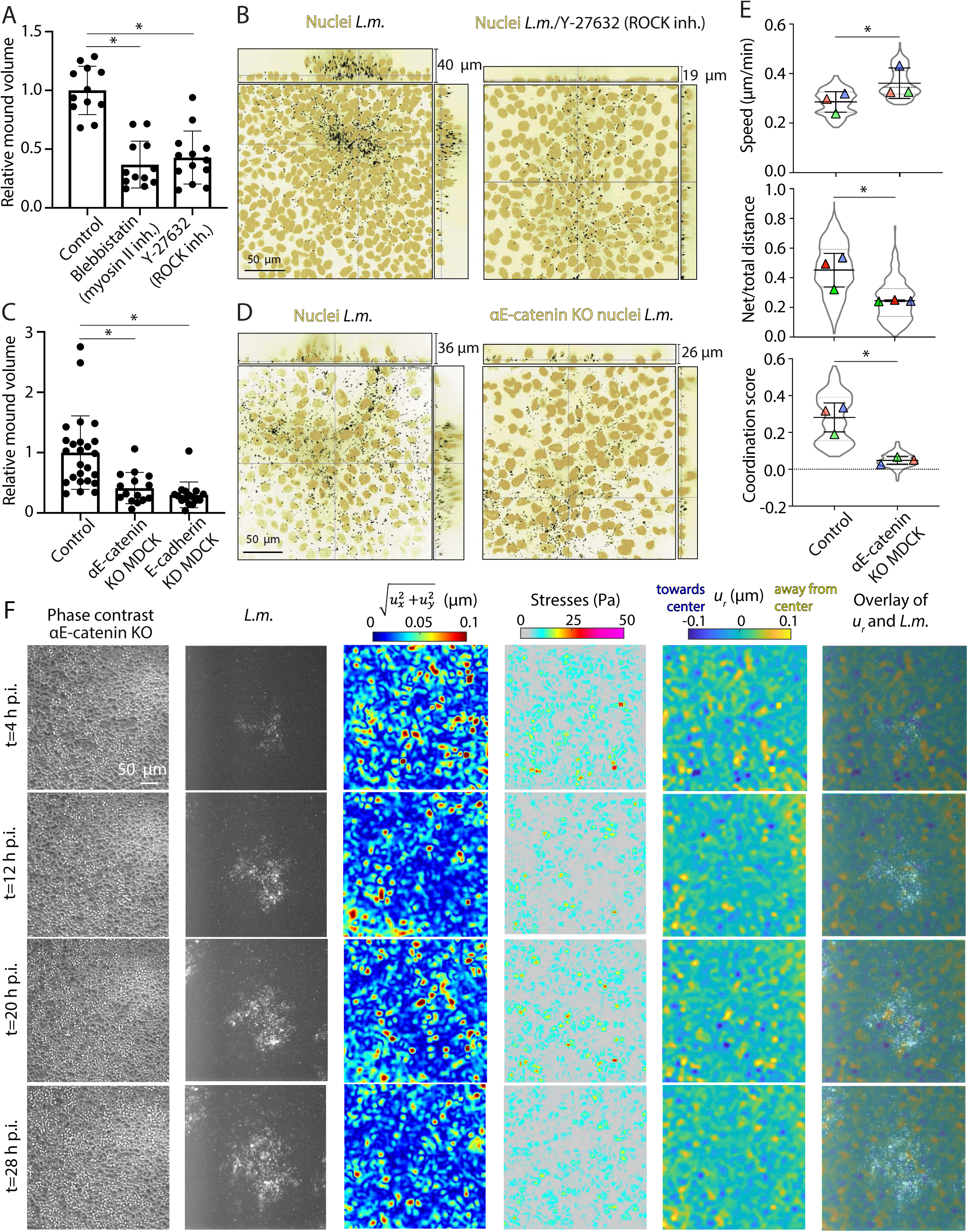
Actomyosin contractility and cell-cell adhesions contribute to mound formation. **(A)** Barplots of relative *L.m.-*infection mound volume at 24 h p.i. for cells treated with vehicle control, 30 µΜ of Y27632, or 50 µΜ of blebbistatin. For each experiment, values have been normalized relative to the mean mound volume of cells treated with vehicle control (mean+/-SD, Wilcoxon Rank Sum test: * p<0.01). **(B)** Representative orthogonal views of MDCK cells treated with vehicle control or 30 µΜ of Y27632 and infected with *L.m.* at 24 p.i. (host nuclei: yellow, *L.m.*: black). **(C)** Barplots of relative *L.m.-*infection mound volume at 24 h p.i. for WT MDCK cells, ΔαE-catenin MDCK or cells treated with siRNA against E-cadherin (siCDH1). For each experiment, values have been normalized relative to the mean mound volume of WT MDCK cells (mean+/-SD, Wilcoxon Rank Sum test: * p<0.01). **(D)** Representative orthogonal views of WT or ΔαE-catenin MDCK cells infected with *L.m.* at 24 h p.i. (host nuclei: yellow, *L.m.*: black). **(E)** Violin plots of mean nuclear speed (µm/min, top), ratio of net to total distance travelled (middle) and coordination score (bottom, see STAR Methods) for WT (n=1072) or αE- catenin knockout MDCK nuclei (n=515). Cells were infected with *L.m.* and imaged between 6-16 h p.i. at 10 min intervals. Dashed line: median, dotted lines: 25 and 75% quartiles. For each experiment (N=3) the average value over the entire field of view and over time is shown as a triangle (mean+/-SD, unpaired T-test: * p<0.01). **(F)** Same as Figure 3C but for αE-catenin knockout MDCK cells infected with *L.m..* See also Figure S4 and Movie S3.

We then examined the role of cell-cell force transduction in eliciting infection mounding by disrupting adherens junctions, using MDCK cells deleted for α-catenin (Ortega et al., 2017) or knocked down for E-cadherin (Figure S4A). In both cases, infection mound formation was dramatically reduced (Figure 4C-D). We went on to examine the kinematics and mechanics of the cells in the infectious foci for αΕ-catenin knockout MDCK cells in more detail. Although αΕ- catenin knockout MDCK move faster than WT MDCK cells between 6-16 h p.i., both their directional persistence and the coordination of migration with their neighbors is significantly lower (Figure 4E). This finding suggests that coordination of movement of neighboring cells though robust cell-cell contacts is necessary for mounding to occur. When we then performed TFM to measure the cell-matrix forces that αΕ-catenin knockout MDCK exert as the infection focus grows, we noted that the overall traction stresses were substantially lower than for WT MDCK cells, and we were unable to observe any differences in the cell-matrix traction stresses of infected mounders versus surrounders (Figure 4F and Movie S3). These observations suggest that cell-cell and cell-matrix forces are tightly coupled, and that a difference in force transduction between the two populations *(i.e.* the infected mounders versus the surrounders) is important for infection mounding to occur.

Interestingly, we found that infection foci for ΔαE-catenin knockout MDCK infected with *L.m.* at 24 h p.i., were much larger than for WT MDCK but also less dense in fluorescence intensity (Figure S4 B-C). This observation is consistent with the idea that mound formation limits the ability of bacteria to spread within the basal cell monolayer. In order to further explore the role of E-cadherin in *L.m.* cell-to-cell spread and formation of infection mounds, we took advantage of the human carcinoma-derived epithelial cell line A431D, which does not express endogenous E-cadherin (Lewis et al., 1997), but which can form very large infection mounds when transduced with human full-length E-cadherin (Figure S1D). The parental E-cadherin null A431D cell line cannot be infected with *L.m.* because E-cadherin serves as the primary receptor for invasion (Ortega et al., 2017), so we mixed A431D cells not expressing E-cadherin with cells expressing full length E-cadherin at a ratio of 100:1 and then infected them with *L.m.*. Under these conditions, bacteria invading the few cells expressing E-cadherin were able to spread into the neighboring E-cadherin null cells, forming large domains of infected cells, but mound formation was dramatically inhibited, similar to our findings with ΔαE-catenin knockout and E-cadherin knockdown MDCK cells (Figure S4D-G). However, the overall size of infection foci for spread into the parental (E-cadherin null) A431D cells was smaller as compared to the control where all A431D cells were expressing E-cadherin, possibly because E-cadherin is directly involved in *L.m.* cell-to-cell spread (Figure S4D), as has previously been shown for *Shigella flexneri* (Sansonetti et al., 1994). Taken together, these findings indicate that cell-matrix and especially cell-cell force transduction during infection is critical for mound formation and for infected cell extrusion.

To examine whether there are any differences in the organization of cell-cell adhesions between infected mounders and surrounders, we performed immunofluorescence for E-cadherin to localize adherens junctions and ZO-1 to localize tight junctions. E-cadherin immunolocalization did not reveal significant differences between infected mounders and surrounders (Figure S2A and S4H). However, ZO-1 localization appeared significantly altered for infected cells, with increased intracellular fluorescence intensity as compared to cells from uninfected wells and surrounders (Figure S4H), consistent with endocytic tight junction remodeling. Next, we explore the relative functional contributions of cell-cell junctions in infected mounders versus surrounders by performing infections in mixed monolayers including both αE- catenin knockout MDCK cells and WT MDCK cells expressing E-cadherin-RFP (to distinguish WT cells from mutants) in varying ratios. When 75% of the host cells were WT, infection mounding occurred normally as for 100% WT monolayers, while when 75% of cells were αE- catenin knockouts, infection mounding was strongly reduced as for 100% αE-catenin knockout monolayers. Intriguingly, when cells were mixed 1:1, we only observed mounding in foci where infected mounders and immediate surrounders happened to be predominately WT in that region of the monolayer, while cells further away from the mound could be of either lineage (Figure S4I). This result suggests that the presence of functional E-cadherin-based cell-cell adherens junctions, at least in the infected mounders and the uninfected cells immediately surrounding the infection mound, is necessary for infection mounding to occur.

### Computational modeling suggests that cellular stiffness, cell contractility and cell-cell adhesions are critical for mounding

Thus far we have shown experimentally that cellular stiffness, cell-matrix and cell-cell force transduction play key roles in the collective extrusion of *L.m*.-infected cells. To better understand how these mechanical signals contribute to infection mounding and explore the precise mechanism of action, we followed a computational approach. We hypothesized that infection mounding could be driven by: (1) decrease in cell stiffness of infected mounders compared to surrounders due to major cytoskeletal alterations; (2) a weakening of cell-matrix contractility of infected mounders making it easier for surrounders to squeeze them; (3) presence of cell-cell adhesions especially for infected mounders and immediate surrounders, allowing the collective squeezing and extrusion of the infection mound; or (4) a combination of the above.

The computational model we built was deliberately simplified, and retained two important features as an initial approach: (1) the problem is formulated in 3D so that changes in cell height could be determined directly; and (2) cells infected with bacteria are permitted to have distinct mechanical properties as compared to uninfected cells, but for simplicity, bacterial infection is considered fixed (*i.e.* bacteria do not spread or replicate). The model assumes that cells form a monolayer, where each cell has a hexagonal shape with six neighbors and can deform in all the directions of the monolayer (Figure 5A). For simplicity of computation, cells are divided in three different parts (Sunyer et al., 2016): the contractile, the adhesive and the expanding/protruding domains (Figure S5B-C). The cells are assumed to have a passive stiffness representing the stiffness arising from the cytoskeleton, including the intermediate filaments, and an active contractility representing the actomyosin contractile apparatus. The interaction of cells with their matrix is also considered through contact interactions and cell-cell adhesions as a continuous material. The assumed mechanotransduction mechanism is depicted in Figure S5A. Briefly, if the cell-matrix displacements are large when cells contract, new cell-ECM adhesions will be formed. If cell-matrix displacements are small and if there is tensional asymmetry, new protrusions will form at the edge of the cells experiencing minimum stress, followed by another round of cell contraction. If there is no tensional asymmetry, there will be no protrusion and the given cell will just contract (Figure S5A). In a stable monolayer at steady state (Figure 5A, case 1), all forces are balanced, and no net cell displacement occurs in each round of the simulation.

**Figure 5.**
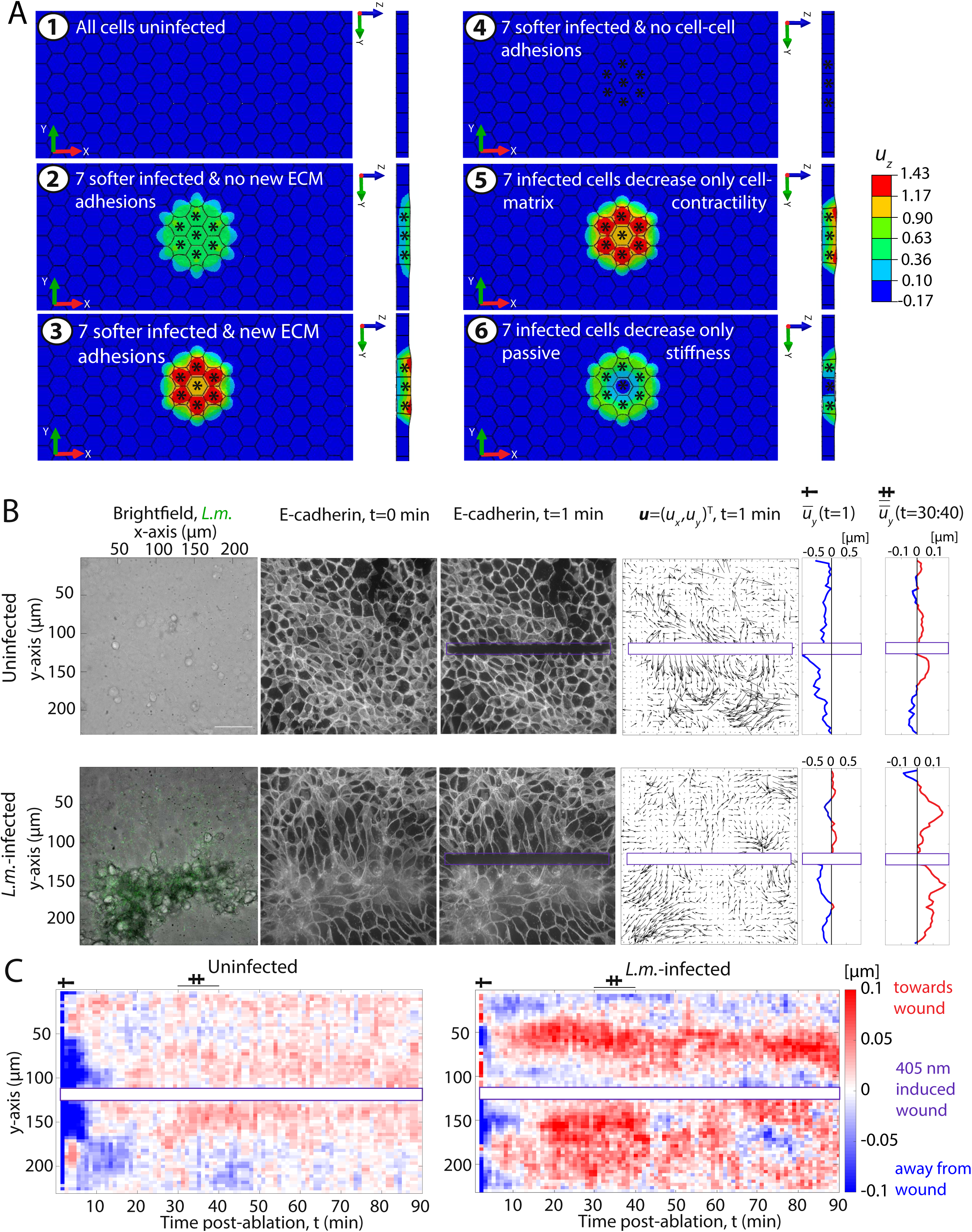
Alterations in cellular stiffness, cell contractility and cell-cell adhesions contribute to mound formation in a computational model, and laser-wounding response is altered in infected monolayers. (**A**) Experiments were performed *in silico*, considering six different cases and assuming that *L.m.* infection leads to softening of infected cells: (1) all cells are uninfected, (2-3) 7 infected cells are softer and uninfected neighbor cells cannot (2) or can (3) create new cell-ECM adhesions, (4) 7 infected cells are softer but all cell-cell adhesions are disrupted in these *in silico* mutants, (5) 7 infected cells are specifically reducing their active contractility and uninfected neighbor cells can form new ECM adhesions, and (6) 7 infected cells are specifically reducing their passive stiffness and uninfected neighbor cells can form new ECM adhesions. Plots show the cell displacements in the vertical (z) direction for the whole model in two different views and considering the six cases mentioned above. Asterisks denote infected cells. (**B**) Laser wounding response of uninfected (top) and *L.m.*-infected MDCK cells in monolayer (bottom) at 24 h p.i.. Columns show: brightfield image of cells superimposed with the *L.m.* fluorescence image (green); corresponding E-cadherin localization prior to wounding; E-cadherin localization at t=1 min post-wounding; total cell displacement vectors **u** (50-fold larger); average vertical displacement u_y_ for t=1 min (positive/negative values point towards/away from the wound and are shown in red/blue respectively); and average vertical displacement u_y_ over the time period between 30 to 40 min. (**C**) Kymographs of the average with respect to the wound displacement vectors, u_y_ as a function of time, t (min, x-axis) and vertical position, y (µm, y-axis) for the same examples as shown in panel B (see single and double crosses). See also Figure S5 and Movie S4.

Using this model, we examined what would happen to the cell displacements and maximum principal stress (maximum tensile stress) in four different cases assuming that infected cells decrease their passive stiffness and their active contractility, as indicated by our experimental results described above. The cases we considered initially were: 1) all cells are uninfected; 2) seven cells are infected; 3) seven cells are infected and uninfected cells in contact with infected ones create new cell-ECM adhesions; and 4) seven cells are infected but cell-cell adhesions are disrupted so no cell-cell force transmission can occur (unlike the other cases in which there will be tight cell-cell contact and significant cell-cell forces). We find that when all cells are uninfected during the contraction phase, cell displacements and principal stresses are symmetrical and cells do not create protrusions, since there is no tensional asymmetry (Figure 5A case 1; see also Figure S5D-F case 1). However, when a cluster of seven cells is infected, we observe that the uninfected cells in close proximity develop tensional asymmetry but since they are unable to form new cell-ECM adhesions, healthy uninfected cells get displaced away from the center of the infection focus (Figure 5A and Figure S5D-F case 2). This outward displacement of surrounding cells is the opposite of our observed experimental result. In marked contrast, if we allow uninfected cells to form new cell-ECM adhesions during the protrusion phase (Figure 5A and Figure S5D-F case 3), we find that uninfected cells in close proximity to infected cells develop tensional asymmetry and are displaced towards the infection focus (Figure 5A and Figure S5D-F case 3). Finally, when cell-cell adhesions are absent, there is no tensional asymmetry and therefore no protrusion formation (Figure 5A and Figure S5D-F case 4). These last two behaviors are consistent with our experimental observations during infection when cells are able to adhere to each other (case 3) or when adherens junctions are compromised (case 4).

Using our model, we can also examine the cell displacements in the vertical direction (z-axis) during *in silico* infection and assess which conditions would give rise to infection mounding. As depicted in Figure 5A where the y-z cross-sections show the vertical displacements of cells in monolayers, the scenario that gives rise to significant initial infection mounding (*i.e.* squeezing of infected cells in the x- and y-axis and an increase in their height in the z-axis) is case 3. This study reinforces the idea that surrounders need to form protrusions towards the infection focus and subsequently new active cell-ECM adhesions are created in order for infection mounding to happen. The orientation and stresses generated by formation of these new adhesions in the *in silico* model are consistent with what we previously observed by TFM (Figure 3C-D and Figure 4E).

Finally, we examined what would happen if only one of the various mechanical cellular parameters were altered during infection, specifically the passive cellular stiffness, or the active cellular contractility, with and without the presence of cell-cell adhesions. If only the active contractility of infected cells decreases to levels lower of those of uninfected surrounder cells, that is sufficient to elicit infection mounding (Figure 5B and Figure S5G case 5). Similarly, if the passive stiffness only of infected cells decreases to levels lower than those of uninfected cells, that is also sufficient to elicit infection mounding, although to a lesser extent (Figure 5B and Figure S5G case 6). Importantly, if passive stiffness of infected cells decreases as compared to uninfected cells but cell-cell adhesions are disrupted, mound formation is abolished completely (Figure 5F and Figure S5C case 4). Altogether, these results indicate that, for infection mounding to occur, it is necessary for infected cells to decrease either their passive stiffness or active contractility (or both), and surrounding uninfected cells need to be able to protrude and form new active cell-ECM adhesions. In addition, cell-cell adhesions are critical since lack of those stalls infection mounding.

### Laser ablation demonstrates alterations in tension at cell-cell junctions for infected epithelial monolayers

Because our computational modeling indicated the importance of tensional asymmetries at cell junctions between infected cells and their uninfected neighbors, we sought to confirm these predictions experimentally. The TFM experiments described above provide direct readouts for cell-matrix force generation, but stresses generated between cells can be inferred only indirectly, and such inference would require assuming that all cells in the monolayer have the same height and stiffness (Serrano et al., 2019). In contrast, laser ablation provides an acute perturbation to cell-cell tension, and has been widely used to determine the direction and relative magnitude of pre-existing stresses in epithelial cell sheets (Fernandez-Gonzalez et al., 2009; Joshi et al., 2010; Kiehart et al., 2000; Kong et al., 2019; Smutny et al., 2015). To determine the nature of changes in epithelial monolayer tension and wound-healing capability on infection, we first ablated rows of cells in uninfected MDCK monolayers and observed a rapid, symmetric recoil of the cells away from the wound on both sides over the first few minutes after wounding, followed by a slower wound closure response that propagated for several tens of micrometers (several cell diameters) from the wound margin and persisted for over an hour (Figure 5C-D). This response is indicative of significant preexisting cell-cell tension throughout the monolayer, followed by post-wounding alterations in cell mechanics near the wound margin concomitant with active crawling to close the wound, and is generally consistent with previous observations on wound response in MDCK cells using other methods for inducing mechanical wounds (Farooqui and Fenteany, 2005; Fenteany et al., 2000; Lv et al., 2020).

Response to laser ablation of cells at the margin of mounds in infected monolayers exhibited several important differences from the uninfected case (Figure 5C-D). First, the magnitude and duration of immediate recoil was substantially less than for uninfected cells, and markedly asymmetric, with the surrounder cells performing less recoil than the infected mounders on the opposite side of the wound. Second, the slower migratory wound-healing response was much more pronounced for both surrounders and infected mounders than for uninfected cells, and extended further away from the wound margin (compare u_y_ over 30-40 min in Figure 5C, and the kymographs for 10 min post-wounding onward in Figure 5D). We used the computational model described above to examine predicted responses given the simple assumptions of that model. Generally, the directions of the predicted recoil (contractile phase) and subsequent migratory response (protrusive phase) for both uninfected and infected monolayers were consistent with our experimental observations (Figure S5H). However, this force-balance model is not well-suited to predict the relative magnitude of longer-term movements because cell migration is explicitly not included in the model.

Overall, we draw two critical conclusions from these experiments. First, the infected mounder cells are clearly capable of mounting a strong wound-healing response, and are able to move away from the center of the infection focus, toward the laser wound margin. This finding reinforces the idea that the cell kinematics associated with mound formation are specifically due to mechanical competition between the surrounders and infected mounders, where the surrounders are stronger, even though the infected mounders retain their normal ability to mount an appropriate directional wound-healing response despite their infected status. Second, both the surrounders and the mounders in the infected condition have significantly altered biomechanics relative to uninfected monolayers, where intercellular tension (driving recoil) is relatively reduced, but the migratory wound response is enhanced.

### RNA sequencing reveals distinct transcriptional profiles among uninfected cells, infected mounders and surrounders

To gain insight into what signaling processes might regulate the mechanical and cytoskeletal changes associated with infection mounding and to understand how those differ in the two battling populations (surrounders versus infected mounders), we followed a transcriptomics approach. To that end, we infected MDCK cells for 24 h with *L.m.* and then used flow cytometry to sort the cells in two groups, infected mounders and surrounders, and extracted their RNA. In parallel, we also extracted the RNA from cells in uninfected monolayers seeded in separate wells at same density (Figure 6A). Through RNA sequencing we found significant numbers of differentially expressed genes (DEGs) when comparing these groups to each other (Figure 6B and Table S1). Principal components analysis (PCA) on these samples revealed three distinct clusters in the PCA space (Figure 6C). PC1 separates cells based on the well they originate from (*i.e.* wells exposed or not exposed to bacteria) whereas PC3 separates cells on the presence of or absence of intracellular bacteria.

**Figure 6.**
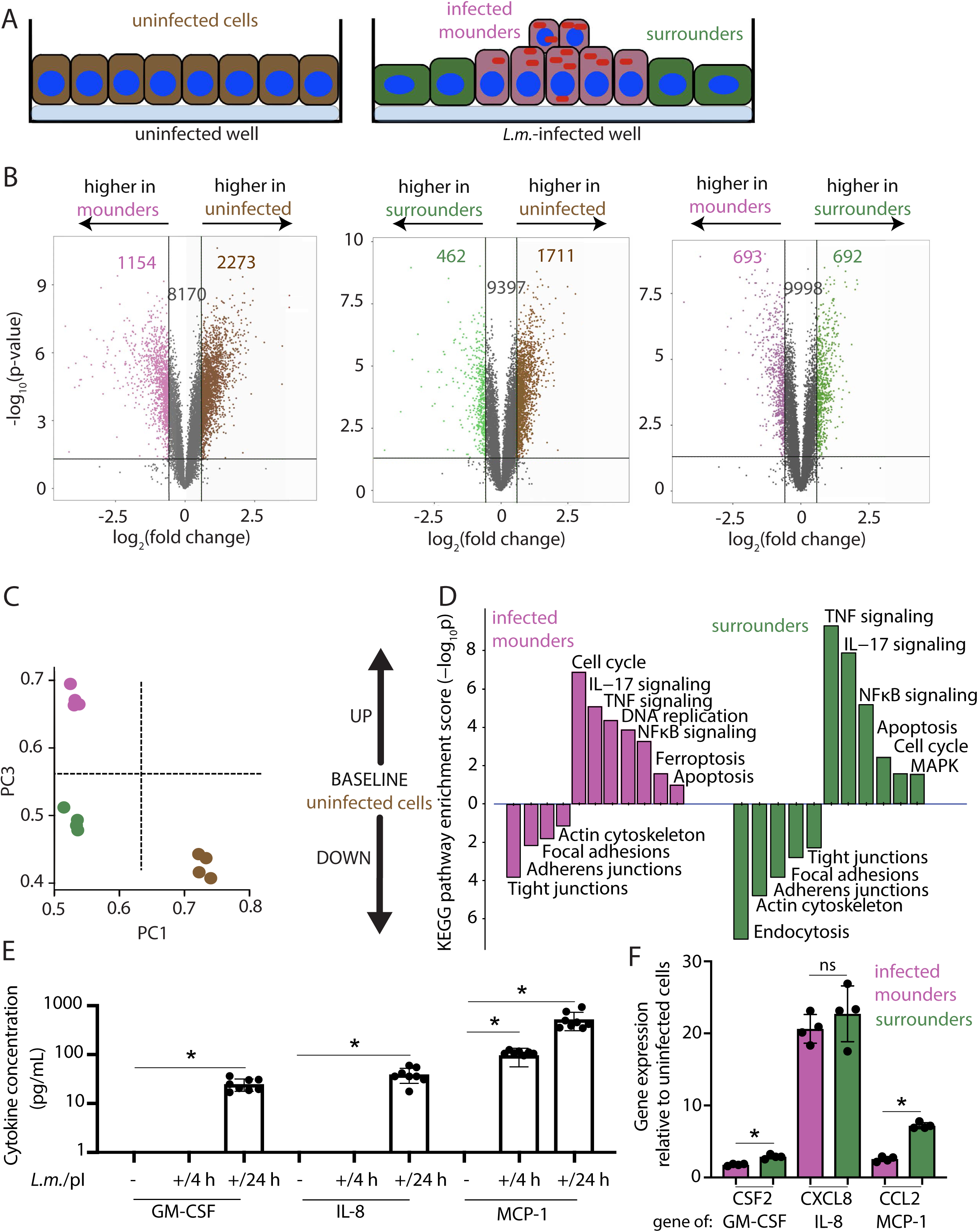
RNA sequencing reveals distinct transcriptional profiles displayed by mounders, surrounders and uninfected cells. **(A)** Sketch showing the different populations that were used for RNA-sequencing (N=4 replicates): uninfected cells from uninfected well (uninfected, brown), infected cells from infected well (infected mounders, pink) and uninfected cells from infected well (surrounders, green). **(B)** Volcano plots showing differentially expressed genes (DEGs) between all groups. The -log_10_ p-values (y-axis) are plotted against the average log_2_ fold changes in expression (x-axis). Non-DEGs are light gray. For each pair of conditions compared upregulated genes of each group are shown in the corresponding color. Number of genes that do not change or are differentially regulated are also shown. **(C)** Principal component analysis (PCA) on top genes that have the ANOVA p value ≤ 0.05 on FPKM abundance estimations. PC1 is plotted versus PC3 for surrounders (green), infected mounders (pink) and uninfected cells (brown). Dashed lines indicate separation based on PC1 on whether cells come from infected or uninfected well and on PC3 based on whether cells are carrying bacteria. **(D)** Pathway enrichment analysis based on the KEGG database on the DEGs identified. Infected mounders (pink) or surrounders (green) are compared to uninfected cells based on their enrichment score (-log_10_p). P-values calculated by Fisher’s exact test were used to estimate the statistical significance of the enrichment of the pathways between the groups compared. **(E)** Barplots showing concentration (pg/mL) of cytokines in the supernatant of cells for cells originating from uninfected well or for cells at 4 or 24 h p.i. (n=8, N=2 experiments, mean+/-SD, Wilcoxon Rank Sum test: * p<0.01). Only cytokines whose expression was >1 pg/mL are displayed. **(F)** Barplots showing gene expression at 24 h p.i. of infected mounders (pink) versus uninfected surrounders (green) as compared to non-infected cells for cytokines encoded by *CSF2*, *CXCL8* and *CCL2* (n=4, mean+/-SD, Wilcoxon Rank Sum test: * p<0.01). See also Figure S6.

We then performed pathway enrichment analysis for the DEGs we identified and discovered multiple pathways significantly perturbed when comparing the different groups (Figure 6D and Table S1). Compared to cells not exposed to infection, infected mounders showed significant downregulation in genes related to tight junctions, with ZO-1 being one of the key genes downregulated. To a lesser degree, groups of genes related to adherens junctions, focal adhesions and regulation of the actin cytoskeleton were also downregulated. Surrounders showed significant downregulation of genes related to endocytosis, the actin cytoskeleton, adherens junctions and to a lesser degree focal adhesions and tight junctions. We also found that many genes whose changes in expression have previously been associated with the epithelial to mesenchymal transition (EMT) were up- or down-regulated for both populations originating from infected wells (infected mounders and surrounders) as compared to uninfected cells that had never been exposed to bacteria. EMT-associated genes that were significantly upregulated included VIM, SNAI1, SNAI2 while genes that were significantly downregulated included ZO-1 and E-cadherin (Table S1). Finally, genes associated with the Rho-family of GTPases crucial for the regulation of the actin cytoskeleton and actomyosin contractility, as well as integrin-related genes, were differentially regulated both for mounders and surrounders as compared to uninfected cells (Table S1). Altogether these results hint at significant infection-related changes in gene expression for regulators of the cytoskeleton, cell-cell and cell-matrix adhesion elements and EMT-associated transcription factors. The observations on upregulation of EMT-associated factors in both surrounders and infected mounders are particularly intriguing in light of their enhanced migratory wound-healing response as compared to uninfected cells, as described above.

Surprisingly, mounders showed upregulation in genes associated with DNA replication and ferroptosis, and both mounders and to a lesser degree surrounders showed upregulation of cell cycle-related and apoptosis pathways. We hypothesized that these findings are hinting at some kind of stress response or death-related process that infected cells are undergoing which might be linked to the mounding phenotype. To test this hypothesis, we first measured the number of cells collectively in uninfected versus infected wells at different MOIs at 24 h p.i.. We found no significant differences in the number of cells between conditions, suggesting that hyperproliferation in infection is probably not occurring (Figure S6A). We then performed the BrdU assay that is commonly used for assaying proliferation (Crane and Bhattacharya, 2013) and also apoptosis (Darzynkiewicz and Zhao, 2011). Only 1% of cells from uninfected wells were BrdU-positive, as compared to 0.5% of surrounders and 8% of mounders (Figure S6B). Interestingly, the mounders that appeared BrdU-positive coincided with the infected cells having the highest *L.m.* fluorescence indicative of the highest bacterial load (Figure S6C), suggesting that BrdU labeling under these conditions most probably reflects apoptosis rather than proliferation. We also performed the TUNEL assay to assess DNA fragmentation in cells from uninfected and *L.m.*-infected wells, and found that the latter exhibited a 2-fold increase in TUNEL-positive cells (Figure S6D). Finally, we used a live-dead stain (Rincón et al., 2018) to identify dead cells after infection, and confirmed through microscopy that many of the extruded cells in the infection mounds do retain the stain (Figure S6E). Overall these findings suggest that cells within in the mounds eventually undergo cell death, reinforcing the idea that infection mounding is a process which is beneficial for the host since infected cells are being cleared out of the monolayer. It also raises the question of whether the infected cells undergo death and are therefore extruded or alternatively if the extrusion of cells from the substrate leads to their subsequent death.

Most importantly, surrounders and to a lesser degree infected mounders showed upregulation of genes associated with the IL-17, NF-κB and TNF innate immune signaling pathways, which indicates that, although surrounders are not themselves infected with bacteria, their distinct mechanical behavior might arise from cytokines released in infection which influence their phenotype, including their gene expression as well as their mechanical behavior. Indeed, when we performed a multiplex immunoassay on the supernatant of cells that were infected for 4 h or 24 h compared to uninfected cells’ supernatant, we found that secretion of several cytokines was enriched during infection (Figure 6E). Out of 12 cytokines and chemokines we tested, we found that GM-CSF (granulocyte-macrophage colony-stimulating factor) was enriched 25-fold, IL-8 was enriched 39-fold and MCP-1 (monocyte chemoattractant protein 1) was enriched 520-fold at 24 h p.i. while the remainder yielded no or low signal (Figure 6E). Interestingly, gene expression of *CSF2* (gene encoding GM-CSF) and of *CCL2* (gene encoding MCP-1) were significantly upregulated to a greater extent for surrounders as compared to infected mounders at 24 h p.i. (Figure 6F and Table S1). Finally, it is of great interest that all three identified cytokines have been previously associated with NF-κΒ signaling, either as being NF-κΒ activators (Ebner et al., 2003; Schreck and Baeuerle, 1990) or as being expressed and secreted in response to NF-κΒ activation (Feng et al., 2006; Hildebrand et al., 2013; Ishikado et al., 2009; Teferedegne et al., 2006) or both (Hoesel and Schmid, 2013; Kang et al., 2007; Manna and Ramesh, 2005). Intriguingly, MCP-1 has also been previously implicated in EMT of cancer cells that adopt a polarized fibroblast-like shape upon exposure to this cytokine (Li et al., 2017).

To assess whether cytokine signaling alone, without actual bacterial infection, can induce the changes in cell polarization and nematic ordering we observed in surrounder cells, we seeded MDCK cells on transwell inserts and infected them (or not) with *L.m.*. Underneath the inserts, we placed cells seeded as monolayers on coverslips so that they would share the same supernatant as the infected cells just above them. When we examined the morphology of MDCK cells on the coverslips 24 h after continuous exposure to supernatant originating from *L.m.-*infected cells, we indeed found significant shape differences as compared to control cells (Figure S6F-H). Cells exposed to infected cell supernatant as compared to control cells had increased aspect ratio and area, and appeared more tortuous in their shape. This result suggests that at least part of the response and collective behavioral change of the surrounder cells originates due to paracrine signaling.

### Inhibition of apoptosis does not attenuate mounding, but perturbations in NF-κΒ signaling do

Given the evidence that infected mounders are undergoing some form of cell death, we sought to understand whether cell death was a cause or a consequence of mound formation. To test whether death of infected cells was required for mound formation, we treated infected monolayers at 4 h p.i. with either the pancaspase inhibitor Z-VAD-FMK or with GSK’872 to inhibit RIPK1 and 3, which are mediators of necroptosis previously associated with *L.m.* infection (Zhang and Balachandran, 2019). We found that infection mounds for cells treated with Z-VAD-FMK and GSK’872 were similar in size to the corresponding mounds of cells treated with vehicle control (Figure 7A-B). This result suggests that cell death occurs after the extrusion of infected cells, not before.

**Figure 7.**
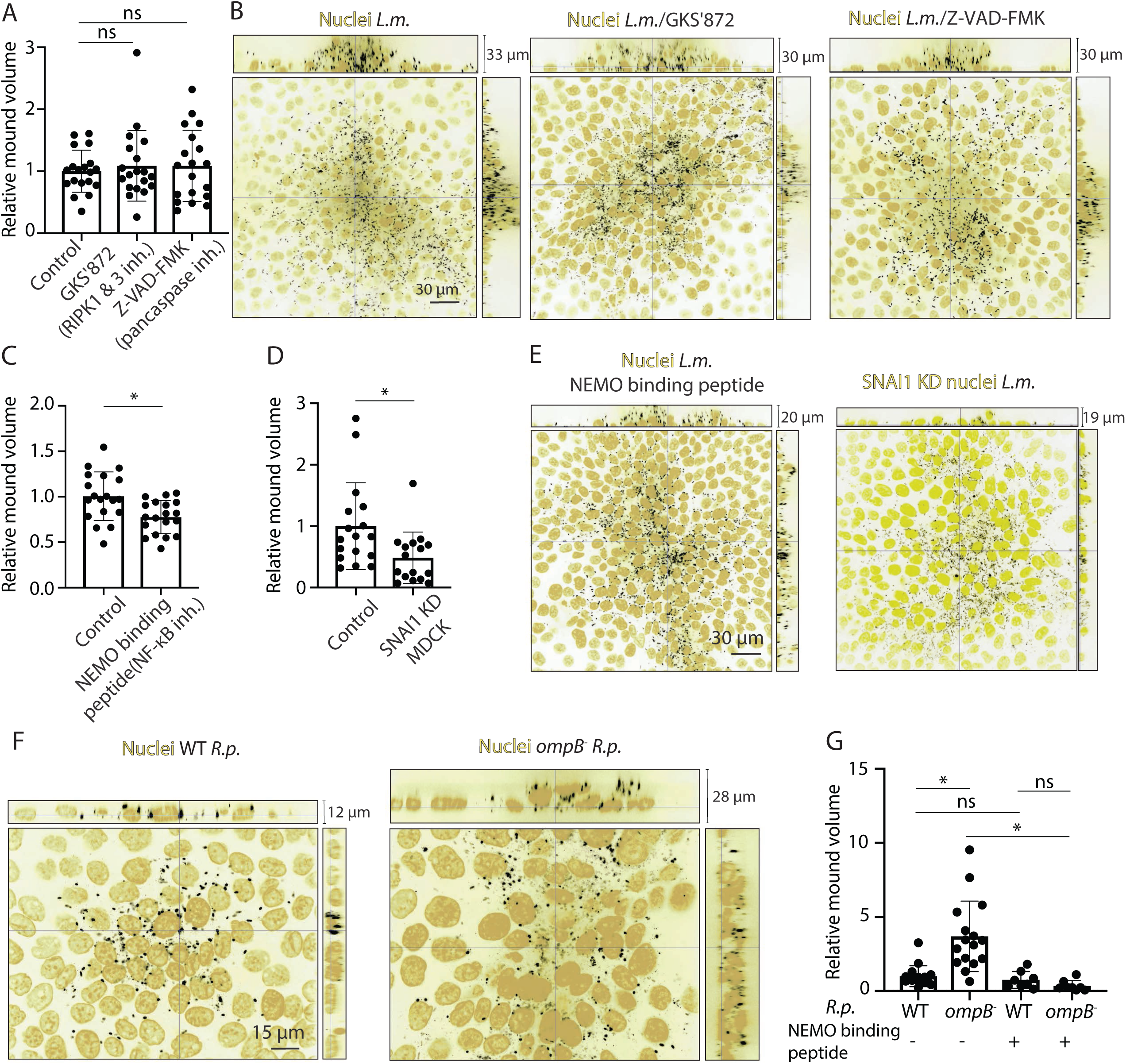
NF-κΒ activation but not cell death contributes to mound formation for two unrelated bacterial pathogens. **(A)** Barplots of relative *L.m.*-infection mound volume at 24 h p.i. for MDCK cells treated with vehicle control, 5 µM GSK’872 (inhibits RIPK1 and RIPK3), or 100 µM Z-VAD-FMK (pancaspase inhibitor). For each experiment, values have been normalized relative to the mean mound volume of cells treated with vehicle control (mean+/-SD, Wilcoxon Rank Sum test: * p<0.01). **(B)** Representative orthogonal views of MDCK cells infected with *L.m.* (host nuclei: yellow, *L.m.*: black) and treated with vehicle control, 5 µM GSK’872, or 100 µM Z-VAD-FMK. **(C)** Barplots similar to panel A showing *L.m.*-infection mound volume at 24 h p.i. for control MDCK cells, or cells treated with 50 µM of NEMO-binding peptide (inhibits NF-κB activation). **(D)** Barplots similar to panel A showing *L.m.*-infection mound volume at 24 h p.i. for MDCK cells treated with control non-targeting siRNA or siRNA against *SNAI1*. **(E)** Representative orthogonal views of MDCK cells infected with *L.m.* and treated with 50 µM of NEMO-binding peptide (left panel) or cells treated with siRNA against *SNAI1* (right panel) (host nuclei: yellow, *L.m.*: black). **(F)** Representative orthogonal views of WT *R.p.-* or *ompB^-^ R.p.*-infected MDCK cells at 72 h p.i. (host nuclei: yellow, *L.m.*: black). P value<0.01 is indicated with an asterisk. See Figure S7. **(G)** Barplots of relative infection mound volume at 72 h p.i. for MDCK cells infected with WT *R.p.* or *ompB^-^ R.p*. (N=2 experiments) and for MDCK cells infected with WT *R.p.* or *ompB^-^ R.p*. and treated with the NEMO binding peptide from 24 to 72 h p.i. (N=1 experiment). For each experiment, values have been normalized relative to the mean mound volume of WT *R.p.-*infected MDCK cells (mean+/-SD, Wilcoxon Rank Sum test: * p<0.01). See also Figure S7.

As described above, we found that NF-κB and NF-κΒ-related pathways (*e.g.* IL-17, TNF) were upregulated for cells originating from infected wells, and that these cells also secreted cytokines related to NF-κB signaling at later times in infection (*e.g.* MCP-1, IL-8). To directly examine whether the transcriptional factor NF-κΒ gets activated late in infection, we performed immunolocalization and indeed observed increased nuclear translocation of NF-κB for cells within infection foci, but not for uninfected cells (Figure S7A-B). In order to determine whether NF-κB translocation in infected cells might contribute to mound formation, we treated *L.m.*- infected cells with the NF-κΒ essential modulator (NEMO) binding peptide that blocks the interaction of NEMO protein with IκB (Dai et al., 2004) therefore inhibiting NF-κΒ activation, which reduces the nuclear:cytoplasmic ratio of NF-κB to background levels (Figure S7B). This treatment with NEMO-binding peptide caused ∼25% reduction in infection mound volume compared to infected cells treated with vehicle control (Figure 7C, E). Treatment of infected cells with the NEMO-binding peptide also suppressed the loss of vimentin from the cells in the mound, to a comparable extent as the loss of cell-cell junctions (Figure S7C-E). This is consistent with the hypothesis that the cytoskeletal changes associated with the decrease in passive stiffness of the infected cells may be downstream consequences of NF-κΒ activation.

Since both MCP-1 and NF-κΒ signaling have been previously implicated in the regulation of EMT genes such as SNAI1 (Li et al., 2017; Pires et al., 2017; Strippoli et al., 2008; Tian et al., 2018; Weichhart et al., 2015) and given the 6- and 9-fold increase in SNAI1 gene expression in surrounders and infected mounders respectively as compared to cells from uninfected cultures, we also knocked down expression of SNAI1 in MDCK cells (Figure S7F), and found a ∼50% decrease in infection mound volume (Figure 7D-E). Overall, these results suggest that innate immune signaling through NF-κΒ and local cytokine signaling play a role in promoting infection mound formation in both the infected mounders and the uninfected surrounders, contributing to the loss of mechanical stiffness in the infected cells and triggering cell shape changes and active motility responses similar to EMT in the uninfected cells immediately surrounding the infection focus.

If our conclusion that NF-κB activation contributes to mound formation is correct, then we hypothesized that we would also observe infection mounding at late times post-infection if we were to infect epithelial host cells with a different intracellular bacterial pathogen that activates NF-κB. To assess this possibility, we infected MDCK epithelial cells with *Rickettsia parkeri* (*R.p.*), an intracellular pathogen unrelated to *L.m.* that is also able to use actin-based motility to drive cell-to-cell spread in epithelial cells (Lamason et al., 2016). We tested two different bacterial strains, WT *R.p.* and the *ompB^-^R.p.* mutant. The outer membrane protein OmpB has been shown to protect the bacterial surface from ubiquitination in the cytoplasm of infected host cells (Engstrom et al., 2019). For *Salmonella enterica* serovar Typhimurium (*S.*Tm.), a intracellular bacterial pathogen that most commonly resides within membrane-bound compartments, bacteria that escape into the cytosol become ubiquitinated, triggering autophagy and activation of NF-κB (Noad et al., 2017). We therefore expected that WT *R.p.* would not trigger significant NF-κB translocation, while the *ompB^-^* mutant would, and confirmed this expectation directly (Figure S7A-B). 72 h p.i., MDCK monolayers infected with *ompB^-^ R.p.* formed large mounds, while those infected with WT *R.p*. did not, and this mounding was abolished by treatment with the NEMO-binding peptide to inhibit NF-κB signaling (Figure 7F-G). Similar to *L.m.*-infected MDCK cells inspected at 24 h p.i., we found that tight junction protein ZO-1 localization was significantly perturbed for mounders in *ompB^-^ R.p.*-infected MDCK cells at 72 h p.i. (Figure S7G). In marked contrast, both ZO-1 localization and the regular spacing of host cell nuclei around infection foci of WT *R.p.-*infected MDCK resembled those of uninfected cells (Figure S7G). Taken together, these results support the idea that the cell mechanical changes associated with mound formation are not simply a result of cytoskeletal alterations driven by pathogens such as *L.m.* and *R.p.* that use actin comet tails for cell-to-cell spread. Rather, NF-κΒ signaling is key to eliciting the mechanical competition between infected mounders and surrounders that leads to mound formation and collective mechanical clearance of infected cells in epithelial monolayers.

## DISCUSSION

Mechanical forces indirectly or directly drive the remodeling of tissues during morphogenesis and are crucial in regulating homeostatic mechanisms such as cell growth, proliferation and death (Ohsawa et al., 2018). In the context of the epithelium, these forces can promote cell extrusion, which is key in preventing accumulation of excess or unfit cells (Grieve and Rabouille, 2014; Gudipaty and Rosenblatt, 2017). Infection of epithelial monolayers with intracellular bacterial pathogens can also trigger mechanical extrusion of individual infected cells, which is typically thought to represent an innate immune protective mechanism on the part of the host (Knodler et al., 2014; Sellin et al., 2014), although in some circumstances it may contribute to bacterial dissemination (Knodler et al., 2010). Here we have described a large-scale, collective mechanical response of epithelial cell monolayers, actively forming infection mounds that can contribute to clearance of domains comprising hundreds of infected cells under conditions where bacterial pathogens can spread non-lytically, such as for *Listeria monocytogenes* (*L.m*.) and *Rickettsia parkeri* (*R.p.*), which are both capable of using actin-based motility for cell-to-cell spread. We argue that the collective infected cell extrusion we describe herein constitutes a process beneficial for the host, since bacteria carried by extruded cells cannot spread along the basal monolayer, limiting the growth of the infection focus. Additionally, extruded cells eventually undergo death, probably due to their forcible separation from the survival signals communicated to normal epithelial cells by direct contact with their basement membrane.

In the geometry associated with the primary site of *L.m.* infection in mammalian hosts, the small intestine, this active extrusion of infected cells driven by changes in the mechanical behavior of the uninfected surrounders would drive the infected cells to be shed into the intestinal lumen, where they could be eliminated from the body by the fecal route. Whether infection mounds would be observed *in vivo* is unclear, since it is possible that physiological shear fluid flows or the presence of immune cells could limit the piling up of infected extruded cells. Intriguingly, bacterial infections in the intestine of *Drosophila melanogaster*, a system that has been widely used as a genetically tractable model for human intestinal infection and pathology (Apidianakis and Rahme, 2011), have been shown to trigger massive remodeling of the intestinal epithelium, featuring both elimination of damaged cells by coordinated delamination, and replacement of the damaged tissue by stem cell proliferation and morphogenetic tissue remodeling (Buchon et al., 2010).

The mechanics of single-cell extrusion have been studied extensively in the context of normal tissue homeostasis as well as in bacterial infection, and mechanisms proposed include lamellipodial protrusions of cells surrounding the extruded cell or a contractile “purse-string” actomyosin ring formed around the extruding cell (Kocgozlu et al., 2016; Pentecost et al., 2006). Contractile “purse-string” mechanisms are also important in the process of wound healing and during development (Abreu-Blanco et al., 2012; Kiehart, 1999). In our system there is no “purse-string” actomyosin ring observed around the infection mound. Instead, mound extrusion depends solely on active crawling on the part of the uninfected cells surrounding the focus of infection, with these surrounding cells becoming morphologically polarized and nematically ordered, generating large outward traction forces on their substrates as they migrate toward the site of infection. Importantly, we have found that both infected cells within the focus (mounders) and their uninfected neighbors (surrounders) undergo a mechanical transition reminiscent of the epithelial to mesenchymal transition (EMT). Both mounders and surrounders exhibit behavioral changes consistent with the most current consensus definitions of the EMT in broad contexts ranging from embryonic development through fibrosis and cancer (Yang et al., 2020). Specifically, both cell populations become less well-ordered, and exhibit increased motility, with strongly reduced intercellular tension and enhanced wound-healing capability as demonstrated by our laser ablation experiments. These diagnostic cell behavioral changes are accompanied by upregulation of canonical EMT-associated molecular markers, most significantly the transcription factors SNAI1 (Snail) and SNAI2 (Slug), and furthermore knockdown of SNAI1 significantly reduced mound formation. Our results suggest that the widely-conserved EMT process can be specifically harnessed in the context of bacterial infection of epithelial cells to ultimately contribute to clearance of infected cells and re-epithelization.

Mechanical competition between two cell populations, such as the ones described herein, where one population loses and gets eliminated and the other wins and eventually gets expanded, have been studied extensively in recent years mostly in the context of oncogenesis (Matamoro-Vidal and Levayer, 2019). Processes leading to the elimination of the “loser” population can include: (1) competition for limiting extracellular pro-survival factors (Senoo-Matsuda and Johnston, 2007); (2) communication (mechanical and/or biochemical) through direct cell-cell contacts (Ohsawa et al., 2018); (3) differential sensitivity to mechanical stress during cell crowding (Hogan et al., 2009; Wagstaff et al., 2016). In the specific case of infection mound formation described here, there is also a clear role for paracrine signaling arising due to infection. Differential sensitivity to mechanical stress between infected mounders and surrounders also plays a key role since both cell stiffness and traction stresses of the two battling populations are dramatically different. Notably, though, the infected mounder cells remain capable of initiating a migratory directional wound-healing response, demonstrating that they are not mechanically completely passive, but rather they are comparatively soft and weak as compared to the stronger surrounders, and cannot resist their coordinated onslaught. Other types of signaling may also contribute to the large-scale coordinated behavior of the surrounder cells. For example, it was recently shown that UV-exposed cells undergo collective cell extrusion over the course of 15 h post-exposure, and that MEK-ERK signaling waves drive this process (Aikin et al., 2019). Whether this type of cell-cell signaling might be involved in infection mounding could be the focus of future studies.

The centrality of NF-κB activation to the innate immune response is evidenced by the wide variety of specific strategies that bacterial pathogens have developed to inhibit or evade it (Rahman and McFadden, 2011; Reddick and Alto, 2014). In this present work, we are particularly intrigued by the central role of NF-κB signaling in formation of infection mounds for both *L.m.* and *R.p.*, because of its well-known involvement in the host’s innate immune response to infection by many bacterial and viral pathogens (Dev et al., 2011). However, it should be noted that NF-κB inhibition attenuates but does not abolish infection mound formation, making it plausible that additional signaling events might contribute to this process. Prior studies have shown that *L.m.* infection of certain host cell types leads to NF-κB activation (Rahman and McFadden, 2011), although the specific trigger(s) from the bacterial side may vary (Drolia et al., 2018; Gouin et al., 2010; Mansell et al., 2000). Interestingly, some non-pathogenic *Rickettsia* bacteria have shown to activate NF-κB while others that are pathogenic dampen the activation of NF-κB (Clifton et al., 1998; Curto et al., 2019; Joshi et al., 2003). In our experiments, we did not observe NF-κB activation in MDCK cells infected with WT *R*.*p.* at 72 h p.i., which correlated with lack of host cell mounding. In contrast, *ompB^-^ R*.*p.* bacteria induced both host cell NF-κB activation and cell mounding. This observation provides one rationale as to why many pathogens that are highly adapted to an intracellular lifestyle, and specifically those that are capable of non-lytic cell-to-cell spread, may have evolved mechanism to suppress or avoid activation of immune signaling pathways that otherwise would lead to mound formation (Rahman and McFadden, 2011; Reddick and Alto, 2014). Furthermore, a large number of viral pathogens are also capable of non-lytic cell-to-cell spread in epithelial tissues (Bird and Kirkegaard, 2015; Sattentau, 2011), and viruses also commonly modulate NF-κB signaling during infection (Zhao et al., 2015). Therefore, we speculate that collective extrusion and mound formation may represent a general innate immune response by host epithelia, which may prove to be relevant for viral as well as bacterial infection.

Our discovery that innate immunity-driven cellular competition promotes the collective extrusion of bacterially*-*infected epithelial cells underscores first, the importance of mechanical signals in regulating infection and second, the relationship between innate immunity signals and cellular mechanics, specifically the EMT. Intriguingly, NF-κB activation has previously been linked to upregulation of genes associated with the EMT in other contexts not involving infection (Strippoli et al., 2008; Tian et al., 2018). Studying the dynamics of these signals and downstream effectors during infection using live-cell biosensors and simultaneously measuring the concurrent mechanical changes will further reveal the crosstalk between innate immunity and cell mechanics and how those in synergy orchestrate the infection process. In this work, we have proposed a novel general cooperative epithelial extrusion mechanism, that could quickly help to limit the local spread of infection in epithelia, particularly in the vulnerable surfaces of the lung and intestine, both common sites of pathogen invasion where the outside world is separated from the inside of the body by a single epithelial monolayer. However, whether these results can be recapitulated *in vivo* in the presence of additional cells (*e.g.* immune cells), varying extracellular matrix topography and presence of external mechanical cues (*e.g.* shear fluid flows) has not yet been explored. Thus, continued investigation using more complex assays will enhance our understanding of the precise biomechanical signals that regulate infection and of how host cells modulate those by orchestrating an innate immune response to clear infection.

## ACKNOWLEDGEMENTS

We are grateful to Nigel Orme and Matthew Footer for preparation of the graphical abstract. We thank M. Footer, M. Walkiewicz, K. Kirkegaard, D. Kiehart and members of the Theriot Lab for discussions and experimental support. We thank A. Hayer for sharing code. We thank the Stanford Cell Sciences Imaging Facility for use of the Atomic Force Microscope. This research was supported by the Cell Analysis Facility Flow Cytometry and Imaging Core in the Department of Immunology at the University of Washington. This part of the project was supported, in part, by Award Number 1S10OD021514-01 from the National Center for Research Resources (NCRR). Its contents are solely the responsibility of the authors and do not necessarily represent the official views of the NCRR or the National Institutes of Health. RNA Seq and RT-qPCR were performed by Arraystar Inc. and multiplex immunoassay by PBL Assay Sciences. This work was supported by NIH R01AI036929 (J.A.T), NIH R01AI109044 (M.D.W), HHMI (J.A.T) and the American Heart Association, Award number: 18CDA34070047 (E.E.B).

## AUTHOR CONTRIBUTIONS

Conceptualization and Methodology, E.E.B., J.A.T., M.D.W, P.E., M.G.B., F.S.A., P.R., J.G.A., M.S.O., J.G.S.; Software, E.E.B, F.S.A., P.R.; Investigation, E.E.B., P.E., M.G.B., F.S.A., P.R., M.S.O.; Writing – Review & Editing, E.E.B., J.A.T., M.D.W, P.E., M.G.B., F.S.A., P.R., J.G.A., M.S.O., J.G.S.; Supervision, J.A.T., J.G.A., M.D.W., J.G.S.

## MATERIALS AVAILABILITY STATEMENT

The RNA sequencing data (FASTq files) generated during this study and subsequent analysis have been submitted to the Gene Expression Omnibus (GEO) database. These data are available at: https://www.ncbi.nlm.nih.gov/geo/query/acc.cgi?acc=GSE140626 and the series record that provides access to all of the data is GSE140626. All the differential expression analysis results of this study are included as supplementary tables in this article.

## STAR METHODS

### Mammalian cell culture conditions

Type II MDCK cells (generous gift from the Bakardjiev lab, University of California, San Francisco) were cultured in high glucose DMEM medium (Thermofisher; 11965092) containing 4.5 g/L glucose and supplemented with 10% fetal bovine serum (GemBio; 900-108) as previously described (Ortega et al., 2017). Passages were between P10-P40. MDCK cells expressing E-cadherin-RFP were a generous gift from the Nelson lab, Stanford University (Perez et al., 2008). αΕ-catenin knockout MDCK cells were also a generous gift from the Nelson lab, Stanford University (Ortega et al., 2017). A431D human epithelial carcinoma cells were generous gift from Cara Gottardi, Northwestern University and were grown in DMEM with high glucose (4.5 g/L) in the presence of 10% FBS and 1% penicillin-streptomycin. Transduced cell lines that express full length E-cadherin were cultured in media supplemented with 800 µg/mL geneticin and were generated as described previously (Ortega et al., 2017).

### Infection of epithelial cells with *Listeria monocytogenes* (*L.m.*)

MDCK or A431D cells seeded on glass or polyacrylamide collagen I-coated substrates were infected as previously described with the following modifications (Bastounis et al., 2018; Ortega et al., 2019). The day prior to infection, host cells were seeded at a density of 2×10^5^ cells/well for cells residing on wells of 24-well plates and grown for 24 h. *L.m.* liquid cultures were started from a plate colony, and grown overnight at room temperature in the dark without shaking, in Brain Heart Infusion (BHI) media (BD; 211059) supplemented with 200 µg/mL streptomycin and 7.5 µg/mL chloramphenicol. Flagellated overnight cultures of bacteria (O.D.600 approximately 0.8) were then washed twice by centrifugation at 2000 x g in PBS to remove any soluble factors. Infections were then performed in normal MDCK growth media by adding 1 mL of bacteria to 15 mL of media and then adding 1 mL of the mixture per well. After 30 min of incubation at 37 °C, samples were washed 3 times in PBS and then media was replaced with media supplemented with 20 µg/mL gentamicin. Multiplicity of infection (MOI) was determined by plating bacteria at different dilutions, on BHI agar plates with 200 µg/mL streptomycin and 7.5 µg/mL chloramphenicol and measuring the number of colonies formed two days post-infection. The resulting MOI was ∼200 bacteria/cell. A similar approach was followed when A431D cells were infected with *L.m.*. The *L.m.* strains used in this study are: JAT607 (Species: *L.m.* 1043S, Genotype/Description: ActAp::mTagRFP) (Ortega et al., 2017), JAT605 (Species: *L.m.* 1043S, Genotype/Description: Constitutive GFP) (Ortega et al., 2017), *ΔinlC* (Species: *L.m.* 1043S, Genotype/Description: *ΔinlC* ActAp::mTagRFP), *ΔinlP* (Species: *L.m.* 1043S, Genotype/Description: *ΔinlP,* obtained from the Bakardjiev lab) (Faralla et al., 2018), JAT983 (Species: *L.m.* 1043S, Genotype/Description: LLOG486D ActAp::mTagRFP) (Rengarajan et al., 2016), *Δhly* (Species: *L.m.* 1043S, Genotype/Description: *Δhly*) (Rengarajan et al., 2016). Flow cytometry experiments of infected host cells were performed 24 h after infection, unless otherwise stated. For drug exposure experiments, unless otherwise indicated, media was removed from the cells and replaced with media containing either the drug or vehicle control 4 h p.i.. Infection mounds were examined at 24 h p.i.. Pharmacological inhibitors used were: para-nitroblebbistatin (target: Myosin II inhibitor, concentration: 50 µM, source: Fisher, NC1706059); Y-27632 (target: ROCK inhibitor, concentration: 30 µM, source: Sigma, 129830-38-2); Z-VAD-FMK (target: pancaspace inhibitor, concentration: 100 µM, source: MilliporeSigma, 5.30389.0001); GSK’872 (target: RIPK1 and RIPK3 inhibitor, concentration: 5 µM, source: Fisher, 64-921-0); NEMO binding peptide (target: NF-κΒ inhibitor, concentration: 50 µM, source: Fisher, 48-003-0500UG).

### Culture and infection of human ileal enteroids with *L.m*

Human ileal enteroids (HIE5) were originally cultured from normal healthy ileal tissue obtained from surgical resection (Holly and Smith, 2018). For continuous culture, human ileal enteroids were maintained in Matrigel domes in human complete crypt culture medium (hCCCM). hCCCM medium is composed of 50% Wnt3a conditioned media (CM), 10% R-spondin-1 CM, 10% Noggin CM, 1X B-27 supplement, 10 mM HEPES, 1X Glutamax, 1X antibiotic-antimycotic, 1 mM N-acetyl-L-cysteine (Sigma), 10 µM Y-27632, 50 ng/ml epidermal growth factor (EGF, Peprotech), 10 nM gastrin (Sigma), 50 ng/ml fibroblast growth factor- 2/basic (Peprotech), 100 ng/ml insulin like growth factor-1 (BioLegend), and 10 mM nicotinamide in DMEM. CM production and quality control were performed as described elsewhere(Holly and Smith, 2018). Medium components are from Thermo Fisher Scientific unless otherwise noted.

Monolayers of human ileal enteroid cells derived from human ileal enteroids were generated as described (Holly and Smith, 2018). Briefly, human ileal enteroids were removed from Matrigel using cell recovery solution (Thermo Fisher Scientific), dissociated in 0.05% trypsin at 37°C for 5.5 min, quenched with DMEM containing 10% FBS, 10 mM HEPES, and 1X Glutamax, and mechanically dissociated with a pipette. Cells were then passed through a 40 µm cell strainer, centrifuged at 400 x g for 5 min and then resuspended in hCCCM. 3 x 10^5^ cells per well were plated onto 96-well thin (∼ 200 µm) polystyrene plates. The wells were first coated with human placental collagen, type IV prior to adding the cells. The media was changed to hCCCM lacking nicotinamide, Y-27632, and antibiotics 1 d after seeding. The cultures were incubated for 5 days prior to *L.m.* infection, with media changes every 2 days. Prior to infection human ileal enteroid cells were incubated with 100 µL of 4 µg/mL Hoechst for 45 min to stain the host cell nuclei. *L.m.* infection was then performed similar to *L.m.* infection of epithelial cells described above. At 24 h p.i. confocal images of *L.m.* infection mounds of human ileal enteroid cells were obtained without fixation since fixation steps detach extruded cells of the infection mounds. We acquired confocal images of the Hoechst-stained host cell nuclei and of the fluorescent bacteria with 0.2 µm spacing using a spinning disk confocal with a 60x 1.4NA Plan Apo oil objective.

### Infection of epithelial cells with *Rickettsia parkeri (R.p.)*

WT *R.p.* strain Portsmouth was originally obtained from C. Paddock (Center for Disease Control and Prevention), and the *ompB^-^* mutant (*ompB^STOP^*::tn) was isolated and validated in a recent study (Engstrom et al, Nat. Microbiol. 2019). *R*.*p*. propagation, purification and infections were carried as previously described (Engstrom et al., 2019). For infection experiments, 3.5×10^5^ MDCK cells were seeded in 24-well plates with sterile circle 12-mm coverslips (Thermo Fisher Scientific, 12-545-80) ∼24 h prior to infection. The following day, 3×10^3^ purified WT or *ompB^STOP^*::tn infectious *R.p.* were used to infect each well in a 24-well plate. Diluted bacterial suspensions were centrifuged onto cells at 300 x g for 5 min at room temperature and subsequently incubated at 33°C for 24, 48, 72 or 96 h as indicated, fixed for 10 min at room temperature in pre-warmed (37°C) 4% paraformaldehyde (Ted Pella Inc., 18505) diluted in PBS, pH 7.4, then washed 2 x with PBS and stored at 4°C prior to immunofluorescence staining.

### Flow cytometry of MDCK cells infected with *L.m*

24 h p.i., infected MDCK cells were detached from the substrate with 200 µL of 0.25% trypsin-EDTA for 10 min. Solutions in each well were pipetted up and down 6 times to ensure single cell suspensions and 200 µL of complete media was added to inactivate trypsin in each well. Suspensions were transferred into 35-µm cell strainers, (Falcon, 352235) and spun through at 500 x g followed by fixation in 1% paraformaldehyde. Samples were then washed once in PBS and stored in PBS with 1% BSA. Flow cytometry analysis was performed on a BD FACS Canto RUO analyzer (University of Washington Cell Analysis Facility). 10,000-20,000 cells were analyzed per each replicate. To ensure analysis of single cells, the bulk of the distribution of cell counts was gated using the forward versus side scatter plot. This gating strategy ensures that single cells are analyzed and debris or cell doublets or triplets are eliminated from the analysis. A second gating step was then performed to exclude cells that autofluorescence by measuring the fluorescence of control-uninfected cells and gating the population of infected cells to exclude autofluorescence.

### MDCK cell transfection with siRNA

For each well of a 24-well plate, 2×10^4^ MDCK cells suspended in serum free media were reverse-transfected with siRNAs at 20 nM final concentration using 0.25 µL lipofectamine RNAiMAX (Invitrogen 13778075). The transfection mix was replaced by full media 8 h later. Synthetic siRNA pools (including 2 distinct siRNA sequences) to target CDH1 and SNAI1 were purchased from Dharmacon (ON-TARGETplus Non-Targeting Pool, catalog number: D-001810-10; Custom CDH1 duplex, catalog number: CTM-521910 and CTM-521911; Custom SNAI1 duplex, catalog number: CTM-544153 and CTM-544153). MDCK cells were treated with control (non-targeting) or experimental siRNA in accordance to the manufacturer’s recommendations. Specifically, to demonstrate that transfection performed was sufficient to get siRNAs into the cells, we transfected cells with synthetic siRNA, siGLO, that makes transfected cells fluorescent 24 h post-transfection To track cell cycle phenotype and to verify that knockdown has occurred with our protocol, we transfected cells with siKif11 which results in substantial cell death of transfected cells approximately 24-48 h post-transfection and can be verified by microscopy using phase optics. Bacterial infections were performed approximately 72 h after transfection.

### RT-PCR

To confirm the knockdowns, MDCK cells treated with control (non-targeting) or experimental siRNA 72 h after transfection (approximate cell concentration was 1.6×10^5^ cells/ well) were harvested and lyzed using the QIAshredder Kit (Qiagen, 79656). mRNA was harvested using the RNeasy Plus Micro Kit (Qiagen, 74004) and eluted in 30 µL RNAase free water RNA concentrations were measured spectrophotometrically (NanoDrop) and were comparable between conditions. cDNA was prepared using the Superscript III First-strand Synthesis SuperMix (Invitrogen, 18080085). RT-qPCR was performed using the SYBR qPCR Master mix by Arraystar Inc. Genes of interest were amplified using the appropriate primers: for CDH1 forward:5’ AGACCCAGTAACTAACGACGG 3’ and reverse:5’ ACACCAAAGTCTTCAGGGATT 3’; for SNAI1 forward:5’ CCCCCATTTGGCTGTGTTG 3’ and reverse:5’ ATCAGTCTGTCGGCTTTTATCCT 3’; for β-actin forward:5’ CCCAGCACAATGAAGATCAAGAT 3’ and reverse:5’ CAAGAAAGGGTGTAACGCAACT 3’. Briefly the steps followed were: (1) perform RT-qPCR for each target gene and the housekeeping gene GAPDH; (2) determine gene concentration using the standard curve with Rotor-Gene Real-Time Analysis Software 6.0; (3) calculate relative amount of the target gene relative to GAPDH.

### Fabrication of polyacrylamide hydrogels and traction force microscopy (TFM)

Polyacrylamide hydrogel fabrication was done as previously described (Bastounis et al., 2018). Glass-bottom plates with 24 wells (MatTek, P24G-1.5-13-F) were incubated for 1 h with 500 µL of 1 M NaOH, then rinsed with distilled water, and incubated with 500 µL of 2% 3- aminopropyltriethoxysilane (Sigma, 919-30-2) in 95% ethanol for 5 min. Following rinsing with water 500 µL of 0.5% glutaraldehyde were added to each well for 30 min. Wells were rinsed with water again and dried at 60°C. To prepare polyacrylamide hydrogels of 3 kPa, mixtures containing 5% acrylamide (Sigma, A4058) and 0.1% bis-acrylamide (Fisher, BP1404-250) were prepared (Bastounis et al., 2018). Two mixtures were prepared, the second of which contained 0.2 µm fluorescent beads at 0.03% (Invitrogen, F8811) for TFM experiments. 0.06% ammonium persulfate and 0.43% TEMED were then added to the first solution to initiate polymerization. First, 3.6 µL of the first mixture without the beads was added at the center of each well, capped with 12-mm untreated circular glass coverslips, and allowed to polymerize for 20 min. After coverslip removal 2.4 µL of the mixture containing tracer beads was added and sandwiched again with a 12-mm untreated circular glass coverslip and allowed to polymerize for 20 min. Next, 50 mM HEPES at pH 7.5 was added to the wells, and coverslips were removed. Hydrogels were UV-sterilized for 1 h and then activated by adding 200 µL of 0.5% weight/volume heterobifunctional cross-linker Sulfo-SANPAH (ProteoChem; c1111) in 1% dimethyl sulfoxide (DMSO) and 50 mM HEPES, pH 7.5, on the upper surface of the hydrogels and exposing them to UV light for 10 min. Hydrogels were washed with 50 mM HEPES at pH 7.5 and were coated with 200 µL of 0.25 mg/ml rat tail collagen I (Sigma-Aldrich; C3867) in 50 mM HEPES at pH 7.5 overnight at room temperature. Next morning, the collagen coated surfaces were washed with HEPES and gels were stored in HEPES.

TFM was performed as previously described (del Álamo et al., 2007; Lamason et al., 2016). Prior to seeding hydrogels were equilibrated with cell media for 30 min at 37°C. Cells were then seeded to a concentration of 2×10^5^ cells per well directly onto the hydrogels 24 h prior to infection. Cells are then either infected or not with low dosage of L.m. and imaging started 4 h post-infection. Multi-channel time-lapse sequences of fluorescence (to image the beads within the upper portion of the hydrogels) and phase contrast images (to image the cells) were acquired using an inverted Nikon Eclipse Ti2 with an EMCCD camera (Andor Technologies) using a 40X 0.60NA Plan Fluor air objective or a 20x 0.75NA Plan Apo air objective and MicroManager software package(Edelstein et al., 2014). The microscope was surrounded by a box type incubator (Haison) maintained at 37°C and 5% CO_2_. Images were acquired every 10 min for approximately 8 h. Subsequently, at each time interval we measured the 2D deformation of the substrate at each point using an image correlation technique similar to particle image velocimetry. We calculated the local deformation vector by performing image correlation between each image and an undeformed reference image which we acquired by adding 10% SDS at the end of each recording to detach the cells from the hydrogels. We used interrogation windows of 32 x 8 pixels (window size x overlap). Calculations of the two-dimensional traction stresses that cell monolayers exert on the hydrogel are described elsewhere (Bastounis et al., 2014; Lamason et al., 2016).

### Atomic Force Microscopy (AFM) for determination of cell stiffness upon infection

A Bioscope Resolve AFM (Bruker, Santa Barbara) was used for MDCK cell stiffness measurements. The AFM operates with the Nanoscope 9.4 software and the data was analyzed using Nanoscope Analysis 1.9 software. The AFM was integrated with an inverted optical microscope (Zeiss AXIO Observer Z1) to allow for correlation with fluorescence and light microscopy. Fluorescence microscopy was used to position the AFM tip over the MDCK cells that resided on glass substrates and were immersed in PBS and was critical for assessing which cells have internalized fluorescent bacteria.

First, we measured cell stiffness using the PFQNM-LC-A-CAL probe (Bruker) that has a conical shape (15 degrees half-cone angle) terminated with 130 nm diameter sphere, and a spring constant 0.096 nN nm^-1^ (pre-calibrated by manufacturer using Laser Doppler Vibrometer). We performed force-volume mapping using a ramp size of 4 µm, windows of 30 µm x 30 µm, 64 samples per line x 64 lines, ramp rate of 10 Hz, and a trigger force threshold of 600 pN. Since the indentation depth of >0.5 µm far exceeded the diameter of the spherical tip apex (130 nm), data were analyzed using the Sneddon Model on the extend curves (no adhesion). A total of 5 different windows were recorded for each condition.

We also probed cellular stiffness using colloidal probes with 5 µm diameter spheres on a pre-calibrated silicon nitride cantilever purchased from Novascan (PT.Si02.SN.5.CAL). Spring constant was 0.16 Nm^-1^ (pre-calibrated by manufacturer). Force-distance spectroscopy measurements were conducted with a ramp rate 0.2 Hz, trigger force threshold 1 nN, and ramp size 6 µm (to accommodate for increased adhesion due to larger contact area). At each field of view, we selected 4 cells and collected 10 measurements per cell. A total of 4 different fields of view were selected for each condition, leading to 16 cells examined per condition. Data were analyzed using the Hertz Model and the extend curves (no adhesion). Each AFM tip was used only for one experiment and then discarded.

### RNA isolation and RNA sequencing

#### Sample Preparation

G-MDCK cells were cultured in high glucose DMEM medium (Thermofisher, 11965092) containing 4.5 g/L glucose and supplemented with 10% FBS (GemBio, 900-108). To generate confluent cell monolayers, 24-well plates glass-bottom for microscopy were coated with 50 µg/ml rat-tail collagen-I (diluted in 0.2 N acetic acid) for 1 h at 37°C, air-dried for 15 min, and UV-sterilized for 30 min in a BSC. MDCK cells were seeded at a density of 2 × 10^5^ cells/well for 24 h. 24 h post-seeding MDCK cells were exposed to *L.m.* (MOI = 200) for 30 min. After washing out the bacteria three times with PBS, media containing 20 µg/mL gentamycin was added to cells to kill extracellular bacteria. 24 h post-infection cells were trypsinized and either left in tubes or sorted into infected and uninfected populations. All tubes containing cells for RNA extraction were treated in parallel (4 replicates per condition). Cells in solution were spun down and lyzed using the QIAshredder Kit (Qiagen; 79656). mRNA was harvested using the RNeasy Plus Micro Kit (Qiagen; 74004) and eluted in 30 µL RNAase free water. A NanoDrop ND-1000 spectrophotometer was used to determine concentration (abs 260) and purity (abs260/abs230) of total RNA samples. Agarose gel electrophoresis was used to check the integrality of total RNA samples (performed by Arraystar Inc.).

#### Library preparation and RNA sequencing

The total RNA was depleted of rRNAs by Arraystar rRNA Removal Kit. We used Illumina kits for the RNA-seq library preparation, which included procedures of RNA fragmentation, random hexamer primed first strand cDNA synthesis, dUTP based second strand cDNA synthesis, end-repairing, A-tailing, adaptor ligation and library PCR amplification. Finally, the prepared RNA-seq libraries were qualified using Agilent 2100 Bioanalyzer and quantified by qPCR absolute quantification method. The sequencing was performed using Illumina NovaSeq 6000 using the High-Output Kit with 2×150 read length. We had an average of 40 million reads per sample. The DNA fragments in well mixed libraries were denatured with 0.1M NaOH to generate single-stranded DNA molecules, loaded onto channels of the flow cell at 8 pM concentration, and amplified in situ using TruSeq SR Cluster Kit v3-cBot-HS (#GD-401-3001, Illumina). Sequencing was carried out using the Illumina X- ten/NovaSeq according to the manufacturer’s instructions. Sequencing was carried out by running 150 cycles.

#### Transcriptome assembly and differentially expressed gene identification

Raw sequencing data generated from Illumina X-ten/NovaSeq that pass the Illumina chastity filter were used for following analysis. Trimmed reads (trimmed 5’, 3’-adaptor bases) were aligned to the reference genome (CanFam3). Based on alignment statistical analysis (mapp.i.ng ratio, rRNA/mtRNA content, fragment sequence bias), we determined whether the results can be used for subsequent data analysis. To examine the sequencing quality, the quality score plot of each sample was plotted. Quality score Q is logarithmically related to the base calling error probability (P): Q = −10log10(P). For example, Q30 means the incorrect base calling probability to be 0.001 or 99.9% base calling accuracy. After quality control, the fragments were 5’, 3’- adaptor trimmed and filtered ≤ 20 bp reads with Cutadapt software. The trimmed reads were aligned to the reference genome with Hisat 2 software. In a typical experiment, it is possible to align 40 ∼ 90% of the fragments to the reference genome. However, this percentage depends on multiple factors, including sample quality, library quality and sequencing quality. Sequencing reads are classified into the following classes: (1) Mapped : reads aligned to the reference genome (including mRNA, pre-mRNA, poly-A tailed lncRNA and pri-miRNA); (2) mtRNA and rRNA: fragments aligned to rRNA, mtRNA; and (3) Unmapped: Reads that are not aligned.

Differentially expressed genes and differentially expressed transcripts are calculated. The novel genes and transcripts are also predicted. The expression level (FPKM value) of known genes and transcripts were calculated using ballgown through the transcript abundances estimated with StringTie. The number of identified genes and transcripts per group was calculated based on the mean of FPKM in group ≥ 0.5. Fragments per kilobase of transcript per million mapped reads (FPKM) is calculated with the formula: 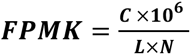, where C is the number of fragments that map to a certain gene/ transcript, L is the length of the gene/transcript in Kb and N is the fragments number that map to all genes/transcripts. Differentially expressed gene and transcript analyses were performed with R package ballgown. Fold change (cutoff 1.5), p-value (≤ 0.05) and FPKM (≥ 0.5 mean in one group) were used for filtering differentially expressed genes and transcripts.

### Principal Component analysis (PCA) and mRNA Function Enrichment Analysis

Principal Component Analysis (PCA) and Hierarchical Clustering scatter plots and volcano plots were calculated for the differentially expressed genes in R or Python environment for statistical computing and graphics. PCA was performed using the plotPCA function in R with genes that have the ANOVA p value ≤ 0.05 on FPKM abundance estimations (Not available for samples with no replicates). Kyoto Encyclopedia of Genes and Genomes (KEGG) pathway analyses of the whole data set of DEG were performed using the R package GAGE “Generally Acceptable Gene set Enrichment” (GAGE v.2.22.0) package implemented in R. The analysis allowed us to determine whether the differentially expressed mRNAs are enriched in certain biological pathways. The p-values calculated by Fisher’s exact test are used to estimate the statistical significance of the enrichment of the pathways between the two groups. The R package “Pathview” v.1.12.0 and KEGGGraph v1.30.0 were used to visualize gene set expression data in the context of functional pathways.

### Immunofluorescence of fixed samples

Uninfected or *L.m.*-infected MDCK cells residing on glass substrates were incubated with 1 µg/mL Hoechst to stain the cells’ nuclei for 10 min at 37 degrees prior to fixation. The following steps were carried out at room temperature. Cells were washed with PBS and fixed with 4% EM grade formaldehyde in PBS for 10 min. Following a wash with PBS, samples were permeabilized for 5 min in 0.2% Triton X-100 in PBS and washed again with PBS. Samples were then blocked for 30 min with 5% BSA in PBS and then incubated with primary antibodies (anti-vimentin: Abcam, ab8069l; anti-pan cytokeratin: ThermoFisher, 53-9003-80; anti-tubulin: Abcam, ab52866; anti-NF-κΒ: Abcam, ab190589; anti-Ε-cadherin: Cell Signaling, 3195; anti-ZO-1: Thermofisher, ZO1-1A12; anti-*R.p.*, I7205 (gift from Dr Ted Hackstadt, NIH Rocky Mountain Laboratories) diluted 1:100 in PBS containing 2% BSA for 1 h for all cases other than anti-*R.p.* where dilution was 1:1000. Samples were washed in PBS three times and then incubated with the appropriate secondary fluorescent antibodies (Invitrogen) diluted 1:250 in PBS containing 2% BSA for 1 h. For actin staining we used 0.2 µM AlexaFluor488 phalloidin (Thermo Fisher, A12379). Samples were washed three times in PBS and stored in 1 mL PBS for imaging. *N* > 8 mounds were analyzed per condition (N> 1200 cells). For epifluorescence imaging, we used an inverted Nikon Eclipse Ti2 with an EMCCD camera (Andor Technologies) and a 40x 0.60NA Plan Fluor air or a 20x 0.75NA Plan Apo air objective and MicroManager software. For confocal imaging we used a Yokogawa W1 Spinning Disk Confocal with Borealis upgrade on a Leica DMi6 inverted microscope with a 50um Disk pattern, a 60x 1.4NA Plan Apo oil objective and MicroManager software.

### Multiplexed magnetic Luminex cytokine immunoassay and data analysis

Samples were centrifuged at 8000 x g to remove debris and assayed immediately. The remaining samples were stored at −80°C. All steps were performed at room temperature. 25 µL of each standard or control was added into the appropriate wells. 25 µL of assay buffer was added for standard 0 (background). 25 µL of assay buffer was added to the sample wells. 25 µL of tissue culture media solution was added to the background, standards, and control wells. 25 µL of sample supernatant was added into the appropriate wells. The beads were vortexed briefly, and 25 µL of the mixed beads was added to each well. We used the Canine Cytokine Magnetic Bead Panel 96-Well Plate Assay (Millipore Sigma, # CCYTMG-90K-PX13). The plate was sealed, wrapped with foil and incubated with agitation on a plate shaker over for 2 h. Following that, the well contents were gently removed, and the plate was washed two times by manually adding and removing 200 µL wash buffer per well. Then, 25 µL of detection antibodies were added into each well. The plate was again sealed and covered with foil and incubated with agitation for 1 h. After incubation, 25 µL Streptavidin-Phycoerythrin was added to each well containing the detection antibodies. (Plate was not washed after incubation with detection antibodies as per manufacturer’s protocol). The plate was sealed and covered with foil and incubated with agitation for 30 min. Then, well contents were gently aspirated out, and the plate was washed two times. Finally, 150 µL of sheath fluid was added to all wells, and the beads were resuspended on a plate shaker for 5 min at room temperature. The plate was run on Bio-Plex 200 flow cytometer. Data analysis was performed using the Bio-Plex Manager 5.0. We used a 5-parameter log fit curve against the signal intensities for the analytes present in the Canine Cytokine Magnetic bead Panel (GM-CSF, IFN-γ, IL-2, IL-6, IL-7, IL-8, IL-15, IP-10, KC-like, IL-10, IL-18, MCP-1, TNF-A) to generate a standard curve for the assay. Sample signal intensities were then back interpolated to the standard curve to determine the analyte concentrations. The upper limit of quantitation (ULOQ) was determined as the highest point at which the interpolated data of the standard curve are within 25% (±) of the expected recovery. ULOQ represents the highest concentration of an analyte that can be accurately quantified. Similarly, the lower limit of quantitation (LLOQ) was determined as the lowest point at which the interpolated data of the standard curve are within 25% (±) of the expected recovery. LLOQ represents the lowest concentration of an analyte that can be accurately quantified.

### Nuclear segmentation, tracking and characterization of dynamics of motion

We acquired time-lapse fluorescence confocal images of Hoechst-stained host epithelial cell nuclei and of the *L.m.* fluorescence using a spinning disk confocal with a 63x 1.2NA Plan Apo water objective at 30 min intervals. For each time point, z stacks including the total height of the cell layer was also acquired with a z spacing of 0.7 µm. To segment and track the host cell nuclei in 3D we used using IMARIS software (Bitplane) by first using the spot detection (segmentation) and then the tracking modules. The x, y and z positions of all identified objects and tracks were exported for import into MATLAB (MathWorks) for further analysis was performed to calculate cell displacements and speeds of migration. For mean square displacement analysis (MSD) we used the @msdanalyzer MATLAB class (Tarantino et al., 2014). We first computed for each cell its MSD as a function of time interval, Δ*t*: *MSD*Δ*t* = 〈|**r***_i_*(*t* + Δ*t*)−**r***_i_*(*t*)|^2^〉, where **r***_i_* (*t*) indicates the position of cell *i* at time *t* and 〈…〉 denotes the average over all time *t*. We then calculated the weighted average of all MSD curves, where weights are taken to be the number of averaged delay in individual curves. In the average MSD plots the grayed area represents the weighted standard deviation over all MSD curves and the errorbars the standard error of the mean. We estimated the self-diffusion coefficient *D*_s_ = lim_Δ*t*→∞_MSD(Δ*t*)/(4Δ*t*) by linear weighted fir of the mean MSD curve on the first 500 min. In the plots shown in Figure 1F-G the grayed area represents the weighted standard deviation over all MSD curves. The error bars show the standard error of the mean which is approximated as the weighted standard deviation divided by the square root of the number of degrees of freedom in the weighted mean. The red line represents the weighted mean MSD curve from which the diffusion coefficient *D* is calculated. Finally, for analysis of the motion type of the objects (i.e. diffusive, subdiffusive or superdiffusive) we performed log-log fitting and modeled the MSD curve by the following power law *MSD(Δt)* = *Г×t*^a^. For purely diffusive motion α=1, for subdiffusive α<1 and for superdiffusive α>1. To determine the power law coefficient we take the logarithm of the power law so that it turns linear: log (*MSD*) = *a*× log(*t*) + log(Г) and by fitting log(MSD) versus log(t) we retrieve α. For more details please refer to @msdanalyzer MATLAB class (Tarantino et al., 2014).

2D tracking of wild-type or αΕ-catenin knockout *L.m.*-infected MDCK cells was performed using epifluorescence microscopy following the movement of Hoechst-stained host cell nuclei between 6 to 16 h p.i.. 2D nuclear segmentation and tracking was performed using IMARIS software (Bitplane). The x, y and z positions of all identified objects and tracks were exported for import into MATLAB (MathWorks) for further analysis of cellular speed, net/total distance travelled and coordination scores as described preveviously (Hayer et al., 2016). Regarding coordination score calculation we considered a radius of 65 µm around each given cell for the calculation of the pairwise velocity correlations (Hayer et al., 2016).

### Cell segmentation and extraction of cell shape characteristics

Epithelial cells were segmented based on pericellular E-cadherin localization using the Tissue Analyzer ImageJ plugin (Aigouy et al., 2016). Masks of individual cells were exported and imported in MATLAB for further analysis. We used custom-made code for calculating the cell area, radial alignment angle θ, aspect ratio and shape parameter 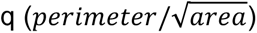. Radial alignment angle θ is defined as the angle between the radial direction and the direction of the major axis of the cell (Figure 2A). If the angle is close to zero, cells are oriented radially towards the center of the image (or the mound). If the orientation is close to 90° the orientation is circumferential. To quantify whether there is any ordering in the orientation of cells, we also calculated for each cell how many of its 100 nearest neighbors share similar orientation angle (Δθ=+/-10^ο^).

### Statistical analysis

For statistical analysis of the kinematics and shape morphometrics of large number of cells characterized in each experiment, we used violin plots considering all nuclei tracked (Figure 1C, H, I and Figure 4E) or all cells characterized (Figure 2D). In the violin plots dashed line represents the median of the distribution and dotted lines the 25 and 75% quartiles considering data from all experiments. For each independent experiment (replicate) the mean value was also calculated and indicated as a colored inverted triangle. Those means when then used to calculate their average (horizontal bar) and standard deviation (vertical bars) and their p-value using an unpaired t-test (Lord et al., 2020).

For characterization of the rest of the data (*e.g.* relative infection mound volumes) data were represented using barplots showing the mean value and standard deviation (vertical bars). P-value was calculated using the non-parametric Wilcoxon Rank Sum test.

### Characterization of mound volume and infection focus area

We acquired confocal images of Hoechst-stained host epithelial cell nuclei and of the fluorescent bacteria in fixed samples using a spinning disk confocal with a 63x 1.2NA Plan Apo water objective. The z spacing used was 0.2 µm for MDCK cells and 0.7 µm for A431D cells, since the field of view imaged for the latter was approximately 9-fold larger. The fields of view were selected so that the infection mounds are approximately at the center of field of view. To quantify the volume of the cells extruded from the monolayer, we first cropped out the lower z slices of each confocal image to exclude cells within the monolayer. We then segmented the channel containing host cell nuclei to form binary images of each mound. An alpha radius of 50 was used to draw a tight boundary around the segmented nuclei using the MATLAB (MathWorks) function alphaShape (Figure S1A-C). The area enclosed within this alpha shape was calculated for each z slice. Lastly, we added up the area of each z slice, converted the area into µm^2^ and multiplied by the increment between each z slice (0.2 µm for MDCK cells and 0.7 µm for A431D cells) to obtain the volume of the mound. An additional step was included in the analysis of αΕ-catenin knockdown MDCK cells and all mounds resulting from infection with *R.p.*. As we observed extrusion of uninfected cells in these cases, we binarized only infected nuclei using input from the bacterial channel. To do so, bacteria were segmented and then enlarged using the imdilate function in MATLAB (MathWorks). This image was applied as a mask to the nuclei channel and only fluorescence within the mask was considered in the analysis. For each different experiment all values were normalized relative to the corresponding values for infected WT MDCK cells treated with vehicle control. Unless indicated three independent experiments were considered and N=13-19 infection mound volumes were calculated in total.

There is no standard metric in the field to measure the efficiency of *L.m.* cell-to-cell spread through a monolayer of host cells. To characterize the area of the infection focus we chose to draw a convex hull, the smallest convex polygon that encompasses a set of points, around the bacteria (Bastounis et al., 2018). This is a computationally inexpensive and consistent way to measure the efficiency of *L.m.* spread.

### Live/dead staining, TUNEL and BrdU assays

We used the NucRed Dead 647 ReadyProbes Reagent (Thermofisher, R37113), a cell impermeant stain that emits bright far-red fluorescence when bound to DNA, for staining dead cells. This reagent stains the cells without plasma membrane integrity and was used to measure cell viability. Briefly, one drop of this reagent was added in each well containing live cells and cells were incubated with this reagent for 20 min before samples were imaged.

In addition, we used the TUNEL assay for DNA fragmentation (Abcam, ab66108) as a means to detect number of apoptotic cells. For staining cells for DNA fragmentation, we followed the manufacturer’s instructions and quantified fluorescein-labeled DNA by flow cytometry for cells at 24 h post-infection. We used the BrdU staining kit for flow cytometry APC (eBioscience; 8817-6600-42) to identify proliferating cells. However, BrdU can also stain apoptotic cells due to the break-up of their genomic DNA by cellular nucleases. For the BrdU assay we followed the manufacturer’s instructions and incubated cells at 24 h post-infection with 10 µm BrdU for 4 h at 37°C before harvesting the cells and proceeding with the above protocol.

### Laser ablation experiments

Laser ablation experiments were performed on *L.m.*-infected MDCK cells at 24 h p.i. using the Firefly system (UGA-42 Firefly; Rapp OptoElectronic) as per manufacturer’s instructions. The region of ablation was approximately 816 pixels in length and a 405 nm laser traversed this region 270 times for a time period of 1 minute passing through the entire height of the mound. During this time, we acquired confocal images of RFP-E-cadherin and the fluorescent bacteria using a spinning disk confocal with a 60x 1.4NA Plan Apo oil objective to determine relaxation of the system and subsequent wound healing. Images were taken with a z spacing of 0.7 µm before and after laser ablation, where the E-cadherin channel was imaged using the 561 nm laser at full power with an exposure of 300 ms, the bacteria were imaged using the 488 nm laser at full power with an exposure of 20 ms, and the brightfield image was taken with an exposure of 50 ms. The cells were imaged every minute for up to 1.5 h post-ablation.

To analyze the displacements of cells upon laser ablation (recoil or relaxation) and their corresponding wound healing response, we used the maximum intensity projections of E-cadherin images of cells and compared subsequent frames starting before laser ablation and following cells every 1 min up to 1.5 h post-laser ablation using particle image velocimetry (PIV)- like technique (Bastounis et al., 2014). For PIV, we used windows of 36 pixels with a 50% overlap. Using the component of the velocity that is perpendicular to the laser ablated segment (u_y_) we then calculated for each instant of time the integral of integral u_y_ along the x-axis, 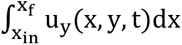. Lining these line integrals over time, we obtained kymographs of the sum of displacements above and below the laser ablated line segment. In these kymographs time, t (min) corresponds to the horizontal axis and position in the y-axis corresponds to the vertical axis (Figure 5D). This representation allows for a detailed quantitative analysis of the recoil behavior of cells upon laser ablation (magnitude of recoil and time it persists) as well as the wound healing response (magnitude, spatial location and temporal evolution).

### Computational modeling of cell monolayers during infection

We assume the cells are arranged in the monolayer as regular hexagons with side length of 7 µm, the thickness of the monolayer is set to 7 µm in the undeformed state following experimental observations and previous computational works (Escribano et al., 2019; O’Dea and King, 2012; Schmedt et al., 2012). We model the mechanical behaviour of the cell distinguishing two main parts: active and passive. The active part represents the actomyosin contractility motors and the passive one the rest of the cytoskeleton (Borau et al., 2011; Moreo et al., 2008). In this way, both parts are assumed linear elastic material, where the total stiffness of each cell is the sum of both (*E_cell_*= *E_active_* + *E_passive_*). Following the experimental AFM measurements, the elastic modulus of uninfected cells is set in 1000 Pa and 250 Pa for the infected cells (*E_cell_*). As a first approach, we assume both parts of the cells are working in parallel, therefore, the active (*E_active_*) and passive (*E_passive_*) elastic modulus are 500 Pa for uninfected cells and 125 Pa for infected cells. We assume the passive part of the cell nearly incompressible, thus the Poisson ratio is set to 0.48 (Moreo et al., 2008). The active part of the cell is mainly actomyosin contraction and we assume this contraction is not isotropic but that it just occurs in the plane of the monolayer. Thus, the Poisson ratio of the active part is assumed zero to uncouple effects in the monolayer plane and in the vertical direction for the active part of the cell.

We model the substrate as a linear elastic material with elastic modulus of 3 kPa corresponding to collagen I-coated polyacrylamide hydrogels used in the TFM experiments. We consider in the model both cell-cell and cell-matrix interactions. We model cell-cell interactions as a continuum and adopt a linear elastic model with stiffness of 4.9 nN/µm and 1.23 nN/µm in normal and shear direction, respectively. When we simulate the inhibition of cell-cell adhesions we set these values close to zero, meaning that cells do not interact anymore mechanically with each other. Meanwhile, the cell-ECM interactions are simulated as cohesive contacts. We assume that cells adhere in different ways to the substrate (Escribano et al., 2018; Sunyer et al., 2016) depending on whether there are cell-cell interactions or not. If the cell-cell adhesions are active (cell monolayer behaves collectively), its adhesion to the substrate is weaker than if the cell-cell adhesions are inhibited (cells forming the monolayer behave individually). The rigid adhesion is assumed to be 1000 nN/µm both in normal and shear direction, and the weak one 10 nN/µm in normal direction and negligible in shear direction. We analyse a short period of time, in which just one contraction and, if it occurs, one protrusion event are simulated. These events could be repeated in time and as a result infection mounding would be more prominent.

To simulate cell ablation, we assume the cell loses its active part just considering the passive part of the cell (Nikolaev et al., 2014; Vileno et al., 2007). Therefore, cell stiffness is reduced and, moreover, ablated cells cannot contract anymore. Cell-cell junctions weaken due to ablation, thus the stiffness of cell-cell junctions in the ablated area is set close to zero during cell contraction and the cell-ECM adhesions are removed (stretching-based interactions). However, during cell protrusion cells are in contact with the ablated area (compression-based contact). We implement the model into a commercial finite element model (FE-based) ABAQUS^70^. To simulate cells two meshes are overlapping sharing the nodes to represent the active and passive components of the cell. We discretise the model with tetrahedral and hexahedral linear elements of average size 2 mm.

## SUPPLEMENTAL INFORMATION

### SUPPLEMENTAL FIGURE LEGENDS

**Supplemental Figure 1.**
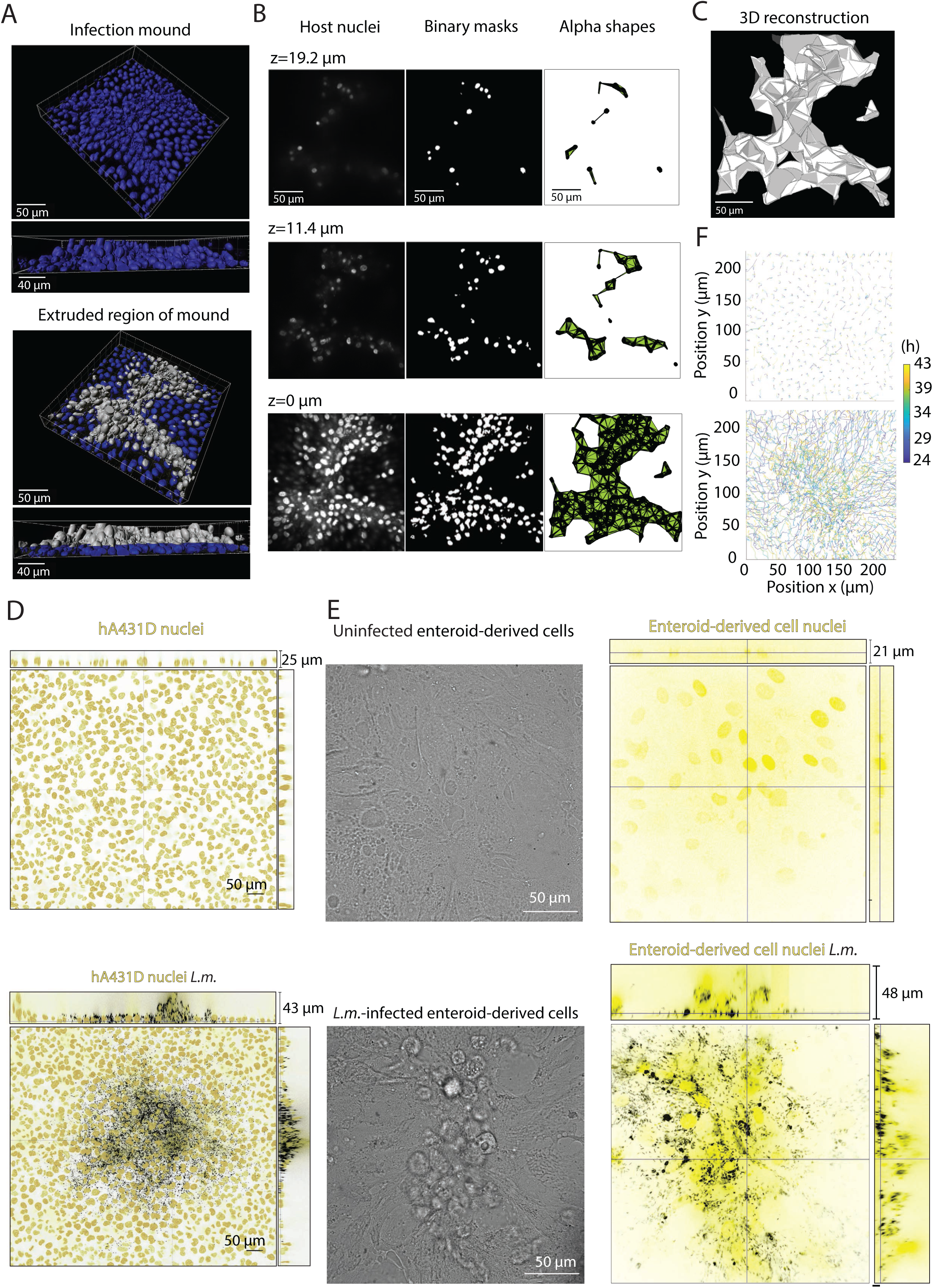
MDCK, human epithelial A431D and ileal enteroid-derived cells infected with *L.m.* form infection mounds at 24 h p.i.. Related to Figure 1. (**A-C**) Depiction of the method followed for extracting the volume of infection mounds. From the confocal image of the region enclosing the mound (A), cells within the monolayer were cropped out, allowing us to calculate the volume of only extruded cells (silver cells in lower images). Each confocal slice was converted into grayscale (representative z slices in B, left), binarized (middle), and contained within an alpha shape (right). By adding up the area of each alpha shape and multiplying by the increment between z slices, we were able to calculate the volume of the mound (reconstructed in 3 dimensions in C). Note that z= 0 µm refers to the first confocal slice corresponding to extruded cells. **(D)** Representative orthogonal views of uninfected (top) and *L.m.*-infected (bottom) human host A431D epithelial cells (host nuclei: yellow, *L.m.*: black). Samples were fixed at 24 h p.i. and on the bottom panel a single infection focus is shown. (**E**) Representative brightfield image of uninfected (top left) and *L.m.*-infected (bottom left) human ileal enteroid-derived cells. Notice the formation of an infection mound in the bottom image. Images on the right show corresponding orthogonal views where host nuclei are in yellow and *L.m.* in black. Samples were imaged at 24 h p.i. (**F**) 2D planar displacements of host MDCK cell nuclei basal layer (*i.e.* of cells attaching to the substratum) for uninfected (top) and *L.m.*-infected MDCK cells tracked between 24 to 44 h p.i.. Nuclear tracks are color-coded based on time.

**Supplemental Figure 2.**
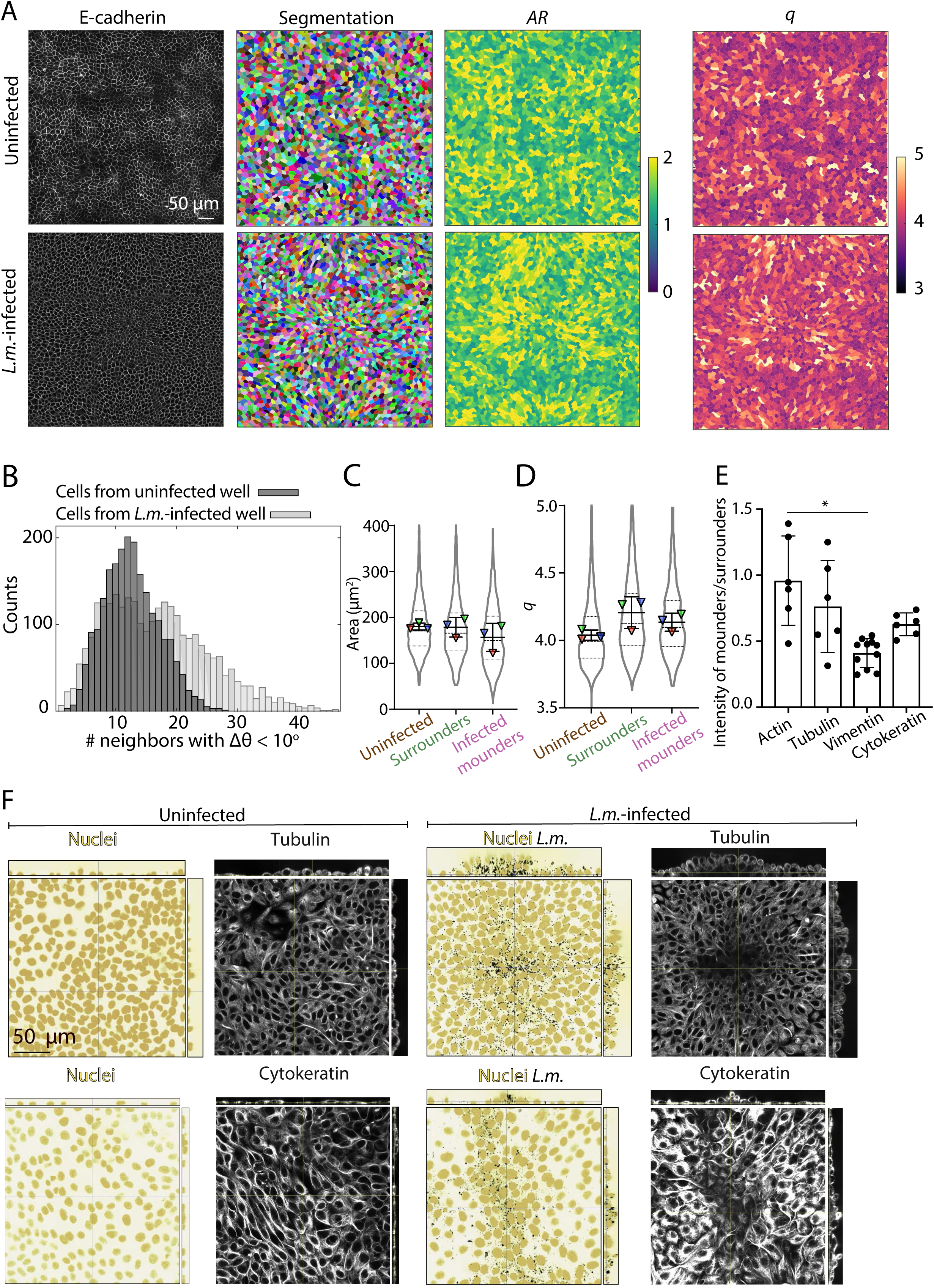
Cell shape, collective behavior and cytoskeletal organization are altered in infected mounders as compared to surrounders. Related to Figure 2. **(A)** Representative images of uninfected MDCK (top) or *L.m.*-infected MDCK (bottom) cells at 24 h p.i., referring to Figure 2A. Columns show: E-cadherin localization used for cell segmentation; cells color-coded based on their aspect ratio (*AR,* major axis/minor axis); and cells color-coded based on the shape factor 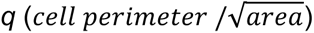. **(B)** Histogram of the average number of neighbors that a particular cell has, which have a similar orientation of their major axis (Δθ<10^ο^). The 100 closest cells based on the nearest neighbor distance of each cell were considered. Dark and light gray histograms correspond to fields of view originating from uninfected and infected wells respectively. (**C-D**) Violin plots of the cell area (µm^2^, C) and of the shape factor *q* (D) for uninfected cells (n=5817), surrounders (n=5163) and infected mounders (n=1248). Dashed line: median, dotted lines: 25 and 75% quartiles. For each experiment (N=3) the average value over the entire field of view is shown as a triangle (mean+/-SD, unpaired t-test: * p<0.05). No significant differences were found. **(E)** Barplot of the per cell ratio of mean fluorescence intensity of actin (N=6 mounds), tubulin (N=6 mounds), vimentin (N=11 mounds) and cytokeratin (N=6 mounds) of infected mounders to surrounders (mean+/-SD, Wilcoxon Rank Sum test: * p<0.01). **(F)** Representative orthogonal views of uninfected (left) and *L.m.*- infected MDCK cells (right) at 24 h p.i.. The image of the host nuclei (yellow) is superimposed to the image of *L.m.* (black) for the infected setting. First and second rows show the corresponding localization of tubulin and cytokeratin respectively.

**Supplemental Figure 3.**
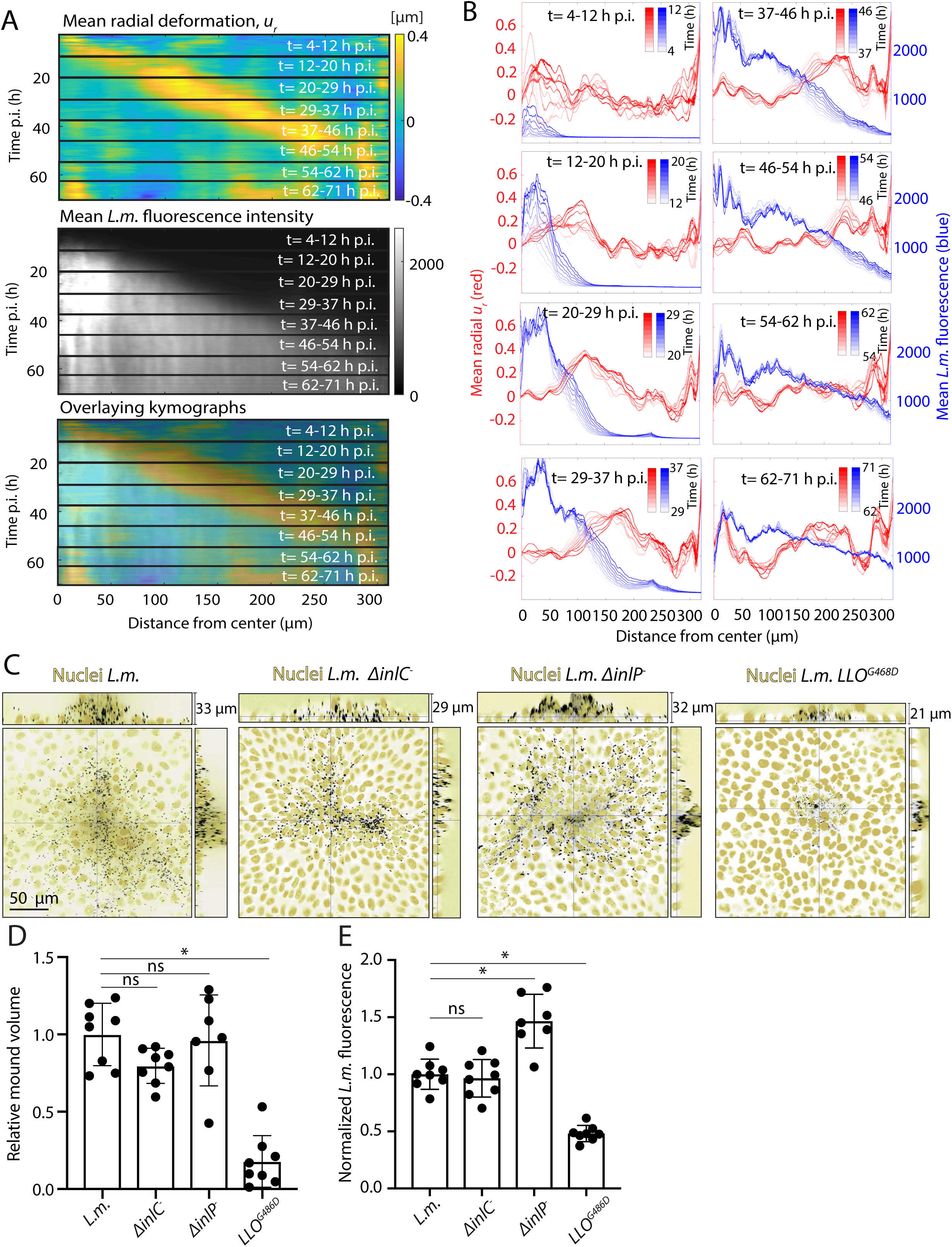
Mound formation requires active traction generation by surrounders, but does not depend on known *L.m.* virulence factors that modify host cell mechanics. Related to Figure 3. **(A)** Kymographs of the mean radial deformation *u_r_* (µm, top), of the mean radial *L.m.* fluorescence intensity (middle) and of their overlay (bottom) as a function of time, t. The center of the polar coordinate system is considered at the center of the infection focus. The kymograph has been split in time segments of approximately 8 h and corresponding time intervals are indicated. **(B)** Mean radial host cell-matrix deformations, *u_r_* (µm) versus distance from the center of the mound outwards is shown in red. Mean radial *L.m.* fluorescence intensity versus distance from the center of the mound outwards is shown in blue. Lines are color coded depending on time p.i. and shown for every 10 frames (50 min). Different plots show different time segments indicated by the corresponding colorbars as infection precedes from 4 h post-infection (top-left) to 62 h post-infection (bottom-right). Panels are chronologically ordered and correspond to the segments of the kymograph shown in panel A. **(C)** Representative orthogonal views of infected host epithelial MDCK with WT *L.m., ΔinlC L.m.*, *ΔinlP L.m.* and *LLO^G486D^ L.m.* (nuclei: yellow, *L.m.*: black). Samples were fixed at 24 h p.i.. **(D)** Barplots showing relative *L.m.*-infection mound volume for MDCK cells infected with WT *L.m.*, *ΔinlC L.m.,ΔinlP L.m.*, or *LLO^G486D^ L.m.* at 24 h p.i.. For this experiment values have been normalized relative to the mean mound volume of cells infected with WT *L.m.* (mean+/-SD, Wilcoxon Rank Sum test: * p<0.01). **(E)** Boxplots showing the relative *L.m.* fluorescence intensity in the infection mounds with respect to mounds of MDCK infected with WT *L.m.*. Infection mounds resulting by infecting either with WT *L.m.,* or *ΔinlC L.m.,* or *ΔinlP L.m.*, or *LLO^G486D^ L.m.* were considered. Samples were fixed at 24 h p.i.. *L.m.* fluorescence was calculated by integrating fluorescence intensity along all 3 dimensions (mean+/-SD, Wilcoxon Rank Sum test: * p<0.01).

**Supplemental Figure 4.**
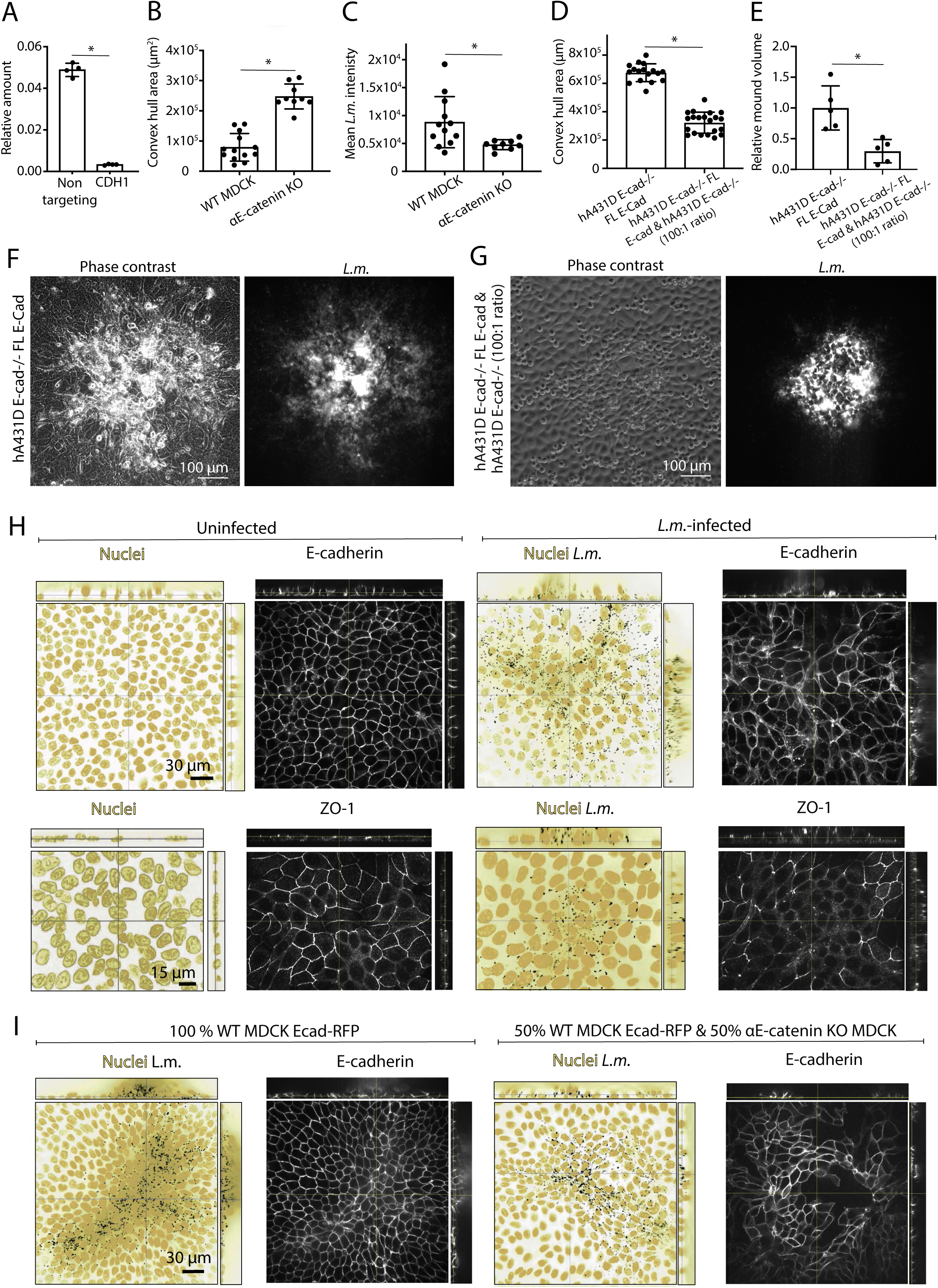
Changes in cell-cell adhesions drive infection mounding. Related to Figure 4. **(A)** Barplots of relative gene expression (normalized to β-actin) of CDH1 (E-cadherin gene) obtained by RT-qPCR for cells treated with control siNT and with siCDH1 (N=4). **(B)** Barplots of the convex hull area (µm^2^) of *L.m.* infection foci for WT or αΕ-catenin KO MDCK host cells at 24 h p.i.. **(C)** Barplots of the mean *L.m.* fluorescence intensity (ratio of the integral of *L.m.* fluorescence intensity to the convex hull area (µm^2^)) for WT or αΕ-catenin KO MDCK infection foci at 24 h p.i.. **(D)** Barplots of the convex hull area (µm^2^) of *L.m.* infection foci for A431D host cells expressing full length E-cadherin or A431D host cells not expressing E-cadherin mixed with cells expressing full length E-cadherin to a ratio of 100:1. **(E)** Barplots showing relative *L.m.*-infection mound volume at 24 h p.i. for A431D host cells expressing full length E-cadherin or A431D cells not expressing E-cadherin mixed with cells expressing full length E-cadherin to a ratio of 100:1. Values have been normalized relative to the mean mound volume for A431D host cells expressing full length E-cadherin. **(F)** Phase contrast image (left) for A431D host cells expressing full length E-cadherin and infected with *L.m.* and corresponding *L.m.* fluorescence (right) at 24 h p.i.. A single infection focus is shown. **(G)** Same as panel F but for *L.m.*-infected A431D host cells not expressing E-cadherin mixed with cells expressing full length E-cadherin to a ratio of 100:1 at 24 h p.i.. (**H**) Representative orthogonal views of uninfected (left) and *L.m.*-infected MDCK cells (right) at 24 h p.i.. The image of host nuclei (yellow) is superimposed to the fluorescence image of *L.m.* (black) for the infected setting. First and second rows shows the corresponding localization of E-cadherin and of ZO-1 respectively. (**I**) Representative orthogonal views of *L.m.*-infected MDCK cells (host nuclei: yellow, *L.m.*: black) expressing E-cadherin-RFP (left) at 24 h p.i. or for 1:1 mixtures of WT MDCK expressing E-cadherin-RFP and αΕ-catenin KO MDCK (right). Data for (A-E) show mean+/-SD, Wilcoxon Rank Sum test: * p<0.01.

**Supplemental Figure 5.**
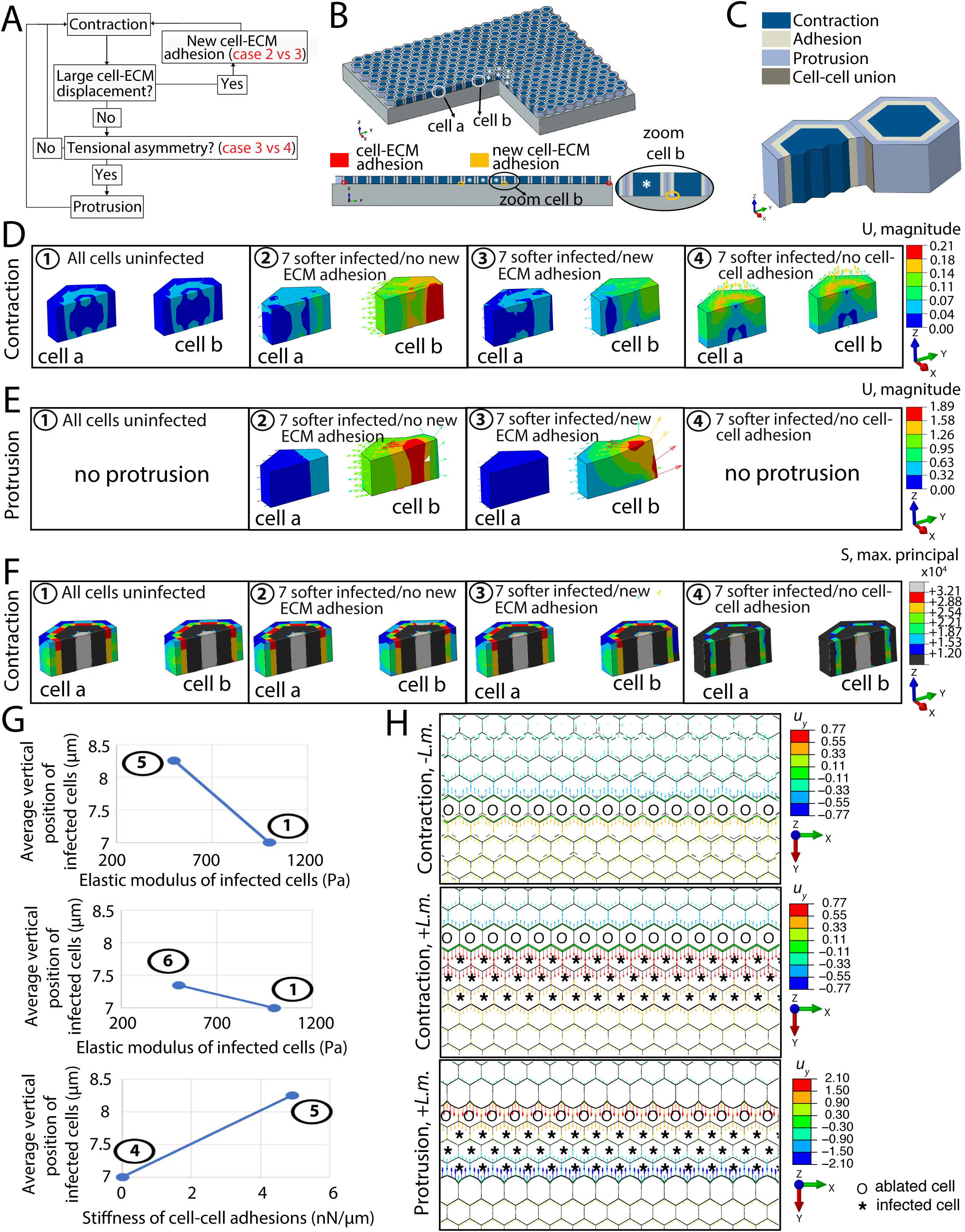
Computational modeling suggests that cellular stiffness, cell contractility and cell-cell adhesions are critical for mounding and wound response. Related to Figure 5. **(A-B)** Schematic showing details of the computational model. Diagram (A) depicts the mechanotransduction mechanism assumed. Schematic (B) illustrates the whole model and indicates the position of two cells that are analyzed below. Infected cells are marked with an asterisk. Underneath of the whole model, sketch shows the two main types of cell-ECM adhesions. The one in red is always present and the yellow only appears in certain cases. **(C)** Sketch showing the three different parts in which cells are divided (contraction, adhesion and expansion) and the cell-cell unions. (**D**) Simulation results showing cell displacements (colorbar shows displacement magnitude in µm and arrows depict displacement direction) during the contraction phase for four different cases assuming that infection leads to softening of infected cells: (1) all cells are uninfected, (2) 7 cells are infected and uninfected neighboring cells cannot create new cell-ECM adhesions, (3) 7 cells are infected and uninfected neighboring cells can create new cell-ECM adhesions and 4) 7 cells are infected but these *in silico* mutants cannot form cell-cell adhesions. (**E**) Same as panel D but showing cell displacements during cell protrusion phase for the four different cases we considered above. (**F**) Same as panel D showing maximum principal stress during contraction in the passive part of the model. Note the stress asymmetry in cases 2, 3 for the cell closer to the infected cell. **(G)** Plots showing the role of the active and passive part of the cell model in driving infection mound formation (see Figure 5A). Note that in case 5 only the active stiffness of infected cells is decreased (*i.e.* the cell contractility) and in case 6 only the passive stiffness of infected cells is decreased (*i.e.* cytoskeletal stiffness). **(H)** Simulation of laser ablation experiments in uninfected (top) and infected monolayers (bottom). Infected cells are indicated with an asterisk, and ablated cells with a circle. Immediately after wound formation, both uninfected and infected cells recoil away from the wound (contraction phase), and then protrude toward the wound site (protrusion phase).

**Supplemental Figure 6.**
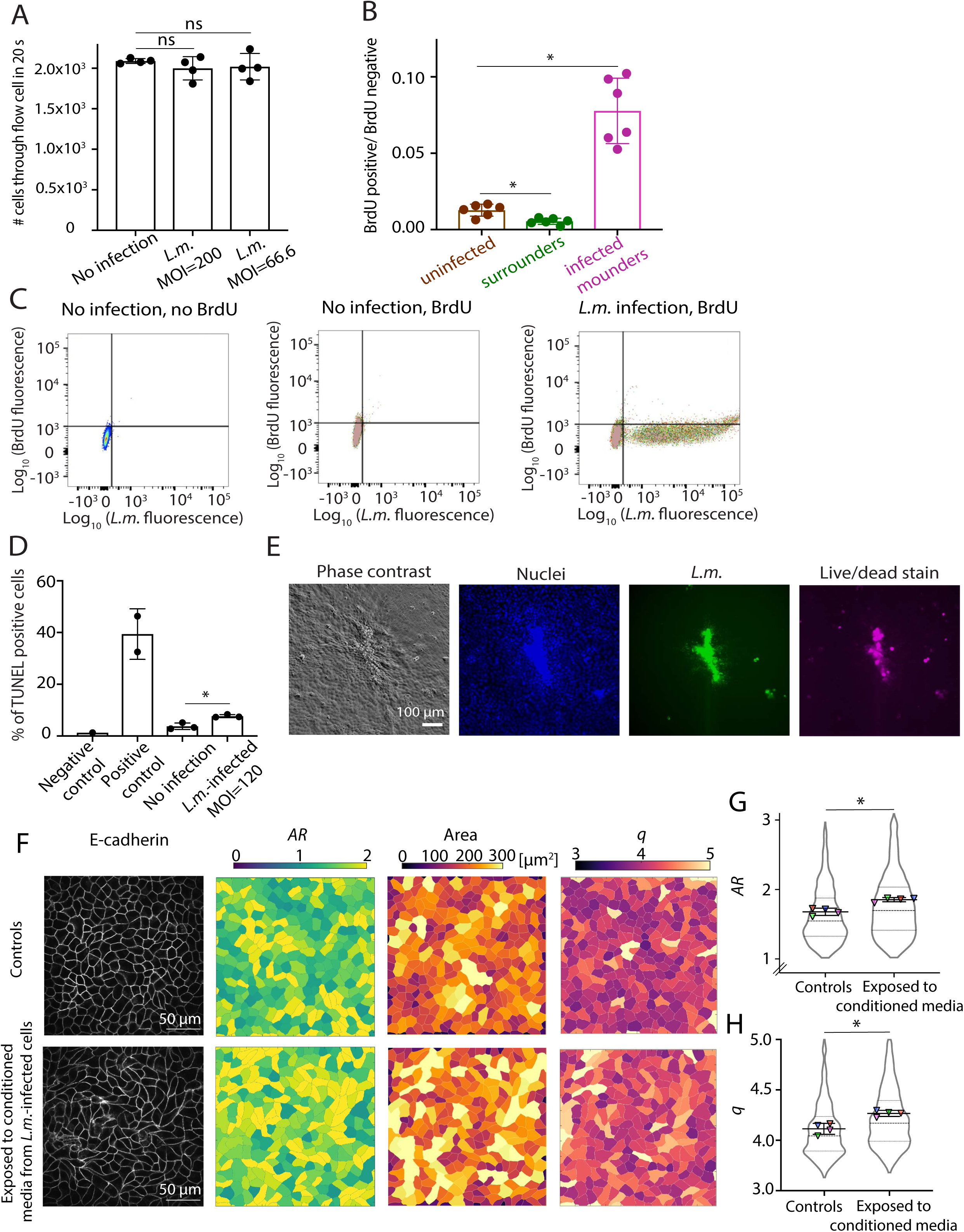
Cytokines released during *L.m.* infection, but not cell death processes, contribute to mound formation. Related to Figure 6. **(A)** Barplot of the number of MDCK cells that pass through the flow cell in 20 s (N=4). Cells originating either from an uninfected or from a *L.m.*-infected well (MOI=200 or 66.66) were collected in flow tubes at 24 h p.i.. Differences in the number of cells were non-significant differences (ns). Data show mean+/-SD, Wilcoxon Rank Sum test: * p<0.01. **(B)** Boxplots showing ratio of BrdU positive to negative MDCK cells for uninfected cells (brown), surrounders (green) and *L.m.*-infected mounders (pink). Cells were stained with BrdU-FITC at 24 h p.i.. *L.m.* expressing mtagRFP was used in infection. Data show mean+/-SD, Wilcoxon Rank Sum test: * p<0.01. **(C)** Bivariate density plot of RFP fluorescence (for *L.m.* containing MDCK cells) versus FITC fluorescence (for BrdU stained cells). Gate separates cells in 4 distinct populations based on whether they are BRdU positive or negative, *L.m.* positive or negative. Left plot shows control uninfected, unstained cells. Middle plot shows uninfected, BrdU-stained cells. Right plot shows *L.m.*-infected, BrdU-stained cells. Examples refer to data shown in panel B. **(D)** Barplot showing the percentage of TUNEL positive cells coming from uninfected (N=3) or *L.m.*-infected wells (N=3). Positive and negative controls supplied with the TUNEL kit used are also shown. Data show mean+/-SD, unpaired t-test: * p<0.01. **(E)** Representative phase contrast images of MDCK infection mounds (1^st^ column), MDCK nuclei (2^nd^ column), *L.m.* fluorescence image (3^rd^ column), and dead cells stained with live-dead stain (4^th^ column). **(F)** Changes in cell shape morphology for cells exposed for 24 h to *L.m.*-infected cell conditioned-media. First row refers to control cells and second row to cells exposed for 24 h to conditioned media. Cells were seeded on transwell inserts and infected or not with *L.m.*. Non-infected cells seeded on coverslips were then placed under the transwell inserts to share the same media as cells seeded on the transwell inserts. 24 h p.i. non-infected cells were fixed, immunostained and inspected via confocal microscopy. Representative image showing E-cadherin localization (1^st^ column), segmented cells colorcoded based on their aspect ratio *AR* (2^nd^ column), on their area (3^rd^ column) and on their shape factor *q* (4^th^ column). **(G-H**) Violin plots of average aspect ratio, *AR* (G) and shape factor, *q* (H) for cells not exposed (n=1059 cells) or exposed to conditioned-media for 24 h (n=924 cells). Dashed line: median, dotted lines: 25 and 75% quartiles. For each replicate (N=4) the average value over the entire field of view is shown as a triangle (mean+/- SD, Wilcoxon Rank Sum: * p<0.05).

**Supplemental Figure 7.**
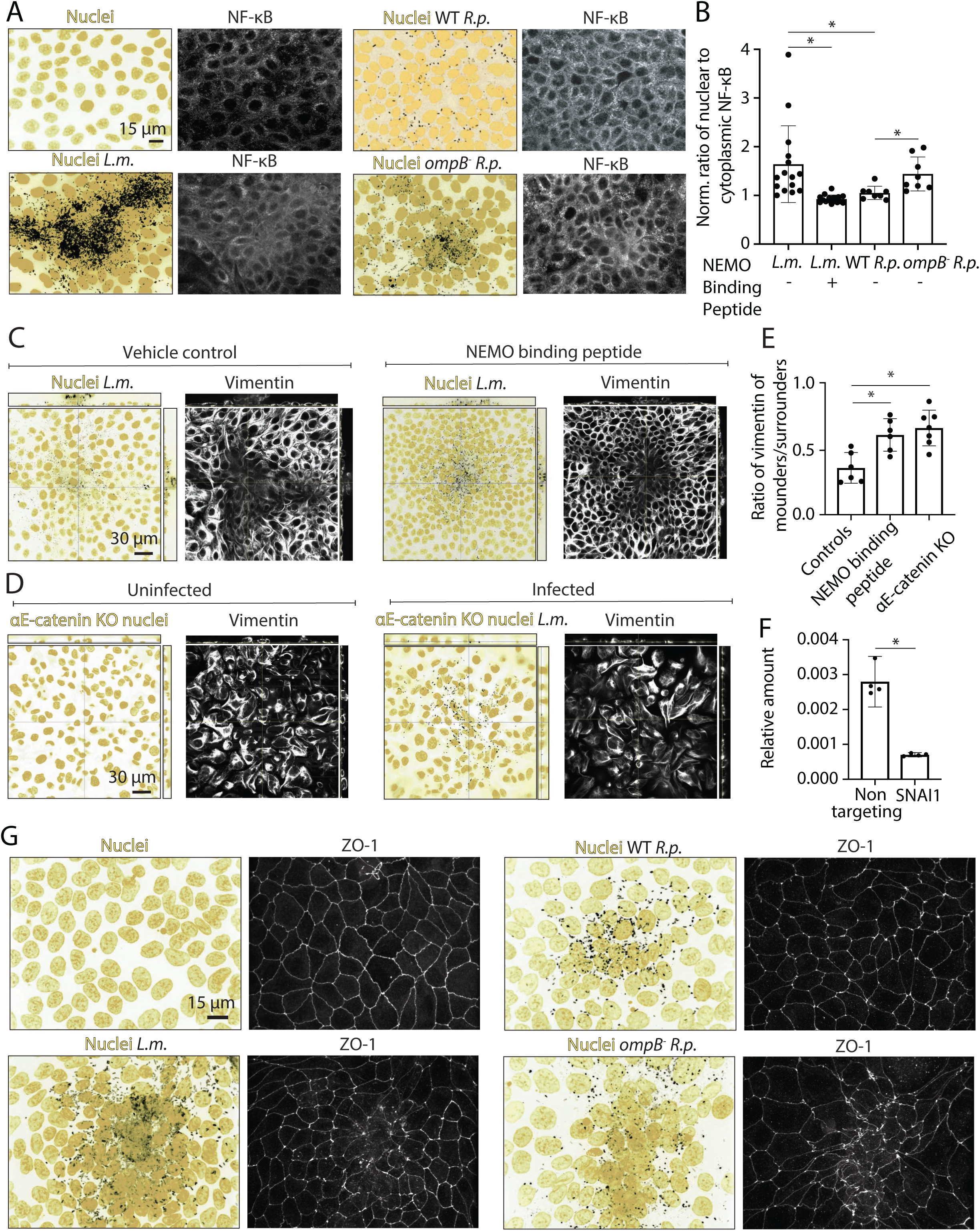
NF-κΒ activation correlates with disruption of vimentin and ZO-1 localization during infection with *L.m.*, WT *R.p.* and *ompB^-^ R.p.*. Related to Figure 7. **(A)** Representative confocal images of the maximum intensity projection of Hoechst-stained host MDCK nuclei superimposed to the maximum intensity projection of bacterial fluorescence around infection foci (left) and of NF-κB localization at a plane located at the basal cell monolayer (right). Uninfected MDCK cells, *L.m.*-infected MDCK cells at 24 h p.i., WT *R.p.*- infected MDCK cells at 72 h p.i., and *ompB^-^ R.p.*-infected MDCK cells at 72 h p.i. are shown. (**B)** Barplots showing the normalized with respect to uninfected cells ratio of nuclear to cytoplasmic NF-κB for *L.m.*-infected MDCK cells treated with vehicle control (24 h p.i.), *L.m.*-infected MDCK cells treated with 50 µM NEMO binding peptide to inhibit NF-κΒ activation, WT *R.p.*-infected MDCK (72 h p.i.) and *ompB^-^ R.p.*-infected MDCK (72 h p.i.). Data show mean+/-SD, Wilcoxon Rank Sum test: * p<0.01**. (C)** Representative orthogonal views of *L.m.*-infected MDCK cells treated with vehicle control (left) or with 50 µΜ of the NEMO binding peptide to inhibit NF-κΒ (right) at 24 h p.i.. The image of the host nuclei (yellow) is superimposed to the fluorescence image of *L.m.* (black) for the infected setting and the corresponding localization of vimentin is also shown. **(D)** Representative orthogonal views of uninfected epithelial αΕ-catenin KO MDCK nuclei (left) and *L.m.*-infected αΕ-catenin KO MDCK cells (right) at 24 h p.i.. The image of the host nuclei (yellow) is superimposed to the fluorescence image of *L.m.* (black) for the infected setting and on the first row the corresponding localization of vimentin is also shown. **(E)** Barplots of ratio of mean vimentin fluorescence intensity per cell of *L.m.*-infected mounders to surrounders (N=6 mounds) for control WT MDCK cells, αΕ-catenin KO MDCK cells and WT MDCK cell treated with the 50 µΜ the NEMO binding peptide to inhibit NF-κΒ. Data show mean+/-SD, Wilcoxon Rank Sum test: * p<0.01. **(F)** Barplots of relative with respect to β-actin gene expression of SNAI1 (Snail gene) obtained by RT-qPCR for cells treated with control siNT and with siSNAI1 (mean+/-SD, Wilcoxon Rank Sum test: * p<0.01). **(G)** Representative confocal images of the maximum intensity projection of Hoechst-stained host MDCK cell nuclei superimposed to the maximum intensity projection of bacterial fluorescence around infection foci (left) and corresponding ZO-1 maximum intensity projection (right). Uninfected MDCK at 72 h post-seeding, *L.m.*-infected MDCK at 24 h p.i., WT *R.p.*-infected MDCK at 72 h p.i., and *ompB^-^ R.p.*-infected MDCK at 72 h p.i. are shown.

## SUPPLEMENTAL TABLE LEGENDS

Table S1. Differentially expressed genes and KEGG pathways between uninfected cells, *L.m.*-infected mounders and uninfected surrounders. Sheets 1-3 of the table present the DEGs identified when comparing the transcriptome of uninfected cells (UU) to surrounders (UI) (1^st^ sheet), uninfected cells (UU) to infected mounders (II) (2^nd^ sheet) and uninfected surrounders (UI) to infected mounders (II) (3^rd^ sheet). Rows correspond to the genes identified and columns A-K correspond to different parameters explained within the table. Columns L-S provide the FPMK of genes in samples for each sample from the corresponding groups compared. Sheets 4-6 of the table show KEGG pathways significantly perturbed when comparing uninfected cells, infected mounders and uninfected surrounders. The gage package was used for pathway analysis which has precompiled databases for mapp.i.ng genes to KEGG pathways so to perform pathway enrichment analysis of DEGs between all three populations. Table shows comparisons performed between uninfected cells (UU) to surrounders (UI) (4^th^ sheet), uninfected cells (UU) to infected mounders (II) (5^th^ sheet) and uninfected surrounders (UI) to infected mounders (II) (5th sheet). KEGG pathways upregulated for each group are indicated. Rows show the KEGG pathways and column different parameters explained in the table including upregulated genes of each category.

## SUPPLEMENTAL MOVIE LEGENDS

**Movie S1. Time-lapse movie showing orthogonal views of uninfected or *L.m.*-infected MDCK host cell nuclei over the course of 20 h. Related to** **Figure 1**. On the left orthogonal views of Hoechst-stained uninfected MDCK nuclei in a confluent monolayer over the course of 20 h are shown, whereas on the right the corresponding views of Hoechst-stained *L.m.*-infected MDCK host cell nuclei over the same time frame are shown. Host nuclei are in yellow and *L.m.* is in black. Acquisition started at 24 h p.i. and frame interval is 30 min. Scale bar and corresponding time p.i. are also indicated. One image from each different plane (*i.e.* x-y, x-z, and y-z planes) is shown.

**Movie S2. Traction force microscopy on WT MDCK cells infected with low multiplicity of *L.m.*. Related to** **Figure 3**. Upper left panel shows the phase contrast image of host MDCK cells. Scale bar and corresponding time p.i. are indicated. Upper middle panel shows the corresponding *L.m.* fluorescence. The infection focus is centered in the middle of the field of view chosen. Upper right panel shows the cell-matrix deformations that cells are producing onto their matrix (color indicates deformation magnitude in µm). Bottom left panel shows the radial deformation, *u_r_* maps (positive deformations values in µm indicate deformations pointing away from the center of the focus, outwards) and bottom middle image the overlap of *u_r_* maps and *L.m.* fluorescence. Bottom right panel shows the traction stresses (color indicates stress magnitude in Pa) exerted by the MDCK host cells that are adhering onto soft 3 kPa hydrogels.

**Movie S3. Traction force microscopy on αΕ-catenin knockout MDCK cells infected with low multiplicity of *L.m.*. Related to** **Figure 4**. Similar to Movie S2 but for αE-catenin knockout MDCK cells.

**Movie S4. Stress distribution during cell contraction and mound formation while uninfected surrounder cells protrude. Related to** **Figure 5**. Simulations showing the stress distribution (mPa) during cell contraction and mound formation (µm) when uninfected cells (surrounders) protrude. Upper left image shows the case where all cells are healthy. Upper right image shows the case where seven cells are infected, and their cell-matrix contractility decreases while uninfected cells in contact with infected ones can create new cell-ECM adhesions. Bottom left image shows the case where seven cells are infected, and their passive stiffness decreases while uninfected cells in contact with infected ones create new cell-ECM adhesions. Finally, the bottom right image shows the case where seven cells are infected, and their passive stiffness decreases but all cell-cell adhesion of both infected and uninfected cells are disrupted.

